# A female-specific CB_1_R-gated subcortical circuit orchestrates defensive homeostasis in risk assessment

**DOI:** 10.1101/2025.09.23.677968

**Authors:** Xue Liu, Hongren Huang, Xiaolong Feng, Xulin Li, Jialin Ye, Huan Yang, Jingjing Liu, Zicong Liang, Zhihan Guo, Ruyi Cai, Shangxuan Cai, Yulong Li, Zhaofa Wu, Liping Wang, Feng Wang

## Abstract

**Highlights:** SC CB_1_R^+^ neurons encode visual survival threats and initiate risk assessment in both sexes

CB_1_R^+^ SC-LHb GABAergic circuit maintains defensive homeostasis in risk assessment in females

eCBs gate this circuit by disinhibiting presynaptic GABA release exclusively in females

Disruption of this mechanism female-specifically impairs risk assessment and stress coping

**In Brief:** Liu et al. identify a female-specific subcortical circuit in which eCB signaling, via CB_1_R on GABAergic terminals in the SC, suppresses transmission to the LHb to maintain precise defensive homeostasis in risk assessment. These findings uncover the circuit and neuronal mechanisms that contribute to the sexually dimorphic maintenance of defensive homeostasis.

Risk assessment in defensive behavior is an adaptive mechanism shaped by natural selection, enabling individuals to evaluate potential threats and thereby maintain defensive homeostasis. However, it remains unknown whether specific neural circuits maintain their behavioral homeostasis in a sex-specific manner. To address this, we investigated its neural basis by hierarchical behavior analysis framework with neural circuit dissection. In mice of both sexes, visual survival threats activated cannabinoid 1 receptor (CB_1_R)-expressing neurons in the superior colliculus (SC) to initiate consistent risk assessment. Deletion of CB_1_R in SC GABAergic neurons female-specifically impairs risk assessment by disinhibiting GABA release in SC-lateral habenula (LHb) projections, resulting in shortened risk assessment. This disruption furthermore increases the occurrence of abnormal spontaneous behavior following chronic stress exclusively in females. We identified a female-specific SC-LHb GABAergic circuit gated by CB_1_R maintains defensive homeostasis in risk assessment. Our findings reveal how a conserved neuromodulatory system sex-specifically gates a subcortical circuit to orchestrate distinct survival strategies.

## INTRODUCTION

Risk assessment during defensive behavior represents an evolutionarily conserved survival mechanism^1–3^, refined through natural selection to optimize trade-offs^3,4^ between immediate survival threats, such as predation risk, and long-term fitness benefits, such as reproductive success. However, how the brain maintains defensive homeostasis in a sex-specific manner remains unknown. Females exhibit heightened risk-assessment capacities^5,6^ to prioritize survival under gestational and lactational constraints^7–9^, manifesting as increased risk avoidance^10^ and perinatal anxiety^11,12^. Conversely, males display stable risk-taking preferences shaped by selective pressures for resource acquisition^13^ and intrasexual competition^14^, where consistent behavioral strategies enhance reproductive success. These sexually dimorphic behavioral strategies, driven by asymmetric reproductive investment^15^ are evident not only in defensive behaviors^16,17^ but also across a range of innate behaviors including mating^18–20^ and aggression^21–23^ to sex recognition^24–26^.

Population viability often depends more critically on female survival in many species^27,28^, Robert Trivers’ theory of parental investment^29^ suggests that females exhibit risk-averse behaviors to protect their greater reproductive investments, a notion supported by our recent findings of sex-specific defensive strategies in mice facing looming threats^30^, suggesting the existence of sexually dimorphic neural mechanisms coordinate by evolutionary pressures. Neural circuits modulating sex-dimorphic innate behaviors frequently involve subcortical hubs that pair conserved anatomy with sex-specific neuromodulation^19,31^. However, it remains unclear whether subcortical structures mediate sex-dimorphic innate defensive behavior, particularly those requiring rapid risk assessment under threat.

The superior colliculus (SC), a conserved hub for processing life-threatening visual cues across vertebrates^32–34^, and the cannabinoid 1 receptor (CB_1_R), known for rapid modulation of survival behaviors^35–37^, emerge as promising candidates. Optogenetic activation of SC parvalbumin^+^ neurons prolongs escape time in female mice^38^, highlighting an evolutionarily driven sexual dimorphism in defensive control. Cannabinoid 1 receptor (CB_1_R), an ancient fear modulator across vertebrates^39,40^, demonstrates sex-specific characteristics in modulating learned fear and fear-related disorders^41,42^, as well as its distribution in the brain^43,44^. It also contributes to the regulation of behaviors essential for survivals, including feeding^35^, sleep^36^, metabolism^37^ and social behavior^45,46^ as well as reproductive strategies, such as maternal behavior^47,48^ and sexual behavior^49^. It is noteworthy that endocannabinoid (eCB), as an endogenous ligand of CB_1_R, achieves synthesis and functional action within milliseconds^50^, matching the temporal demands of life-or-death defensive responses mediated by subcortical circuits such as the SC. We therefore hypothesize that the CB_1_R in SC may gate defensive homeostasis through a sex-specific subcortical fast regulatory mechanism.

Here, we resolve this gap by integrating the hierarchical behavior analysis framework (HBAF)^51^ with neural circuit dissection. We demonstrate that risk assessment in response to visual threats is encoded by SC CB_1_R^+^ neurons in both sexes, while CB_1_R in GABAergic SC-lateral habenula (LHb) projections is necessary for female-specific defensive homeostasis in risk assessment. eCB release from the LHb activates the CB_1_R on the terminal projecting from the SC, while the deletion of CB_1_R in SC GABAergic neurons disinhibits presynaptic GABA release, resulting in a female-specific impairment in risk assessment. Disruption of CB_1_R in SC GABAergic neurons induces female-specific stress vulnerability. Importantly, we identify a neural substrate for evolutionarily optimized sex differences in survival strategies, with female-specific defensive homeostasis in risk assessment directly linked to vertebrate-conserved eCB system. We demonstrate how this circuit integrates evolutionary constraints with real-time threat evaluation and establish a general mechanistic paradigm in which conserved neuromodulatory systems maintain sex-specific defensive homeostasis through subcortical disinhibition.

## RESULTS

### SC CB_1_R^+^ neurons are necessary for processing visual threats in both sexes

To access CB_1_R^+^ neurons genetically, we generated a CB_1_R^iCre^ knock-in mouse line (Figure S1). *In situ* hybridization confirmed >80% of *CB_1_R* mRNA co-localization with Cre protein in both sexes (Figure S1K), exceeded 90% in the superior colliculus (SC) (Figure S1I, K). Quantitative analysis revealed that CB_1_R^+^ cells make up more than 50% of the SC neuronal populations, independent of sex (Figure S2A-B). To investigate activation due to threats, we profiled c-Fos mRNA expression patterns following looming stimuli (Figure S3A). Looming significantly increased c-Fos mRNA expression and its co-localization with Cre proteins in the SC of both sexes (Figure S3B-E). Remarkably, >80% of c-Fos^+^ neurons expressed CB_1_R (Figure S3F), indicating predominant engagement of CB_1_R^+^ neurons in threat processing. Additionally, we monitored real-time SC CB_1_R^+^ neuronal activity via calcium imaging during looming exposure (Figure 1A-B). By injecting either unilateral AAV2/9-EF1α-DIO-GCaMP6s or a control virus into the SC of CB_1_R^iCre^ mice, we enabled cell-specific recordings (Figure 1A). During looming presentation, we observed strong calcium transients in SC CB_1_R^+^ neurons in both male and female mice (Figure 1C-D), indicating sex-conserved recruitment.

**Figure 1.**
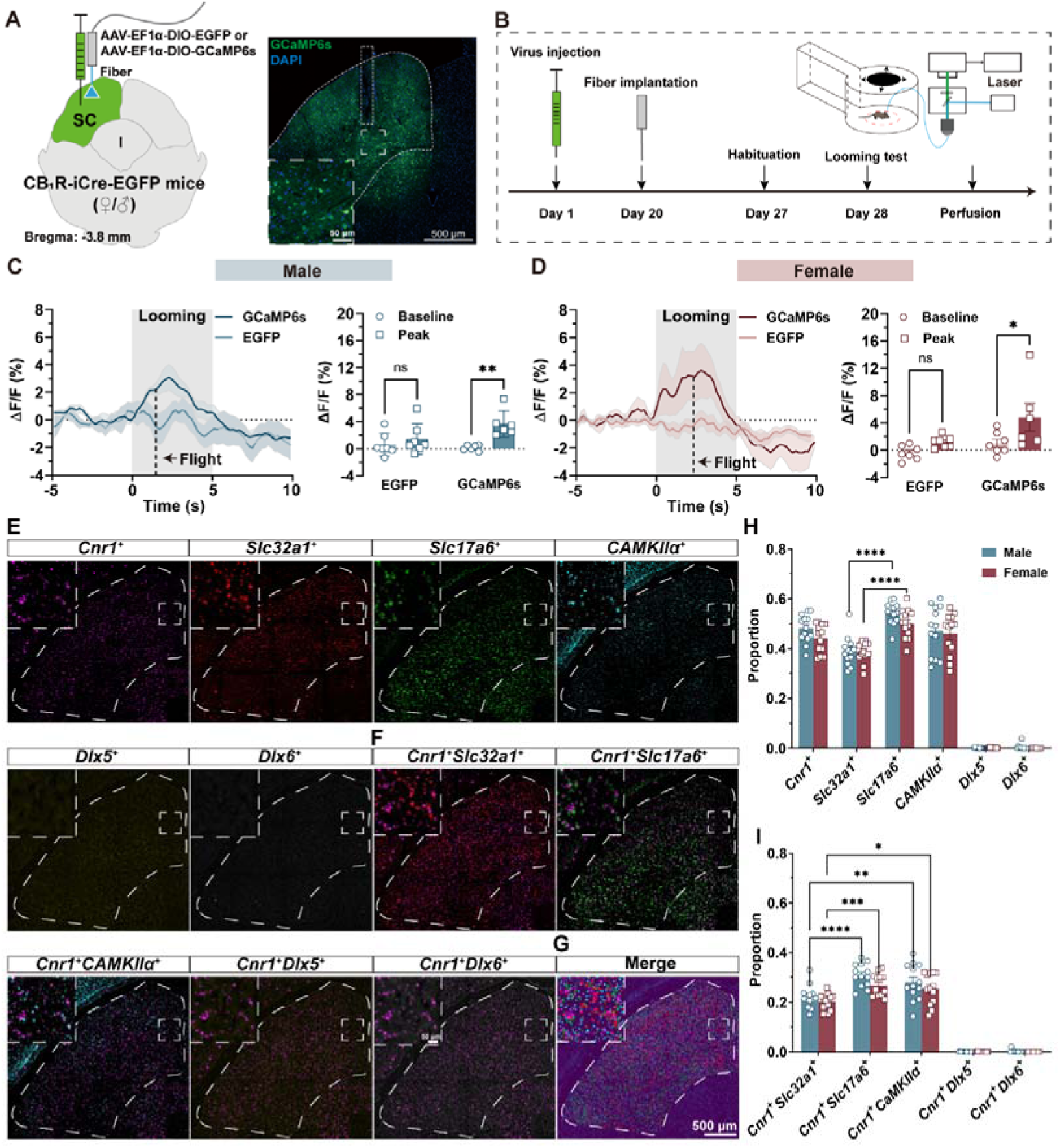
SC CB_1_R^+^ neurons are activated during visually evoked defensive behavior. (A) Left: Schematic representing viral injection of AAV2/9-EF1α-DIO-GCaMP6s or AAV2/9-EF1α-DIO-EGFP into the unilateral SC of CB_1_R-iCre-EGFP mice. Right: Representative image showing the recording site within the SC. Scale bar, 500 μm and 50 μm (enlarged view). (B) Experimental timeline and schematic of the looming stimuli paradigm used to record calcium signals via fiber photometry in a refuge-containing open field apparatus. (C and D) Left: Average calcium transients for the entire test group in males (C) and females (D). Shaded areas represent error bars, while the gray shadow indicates the looming stimuli event. Right: Plots showing calcium transients during looming stimuli in males (C) and females (D). Unpaired Student’s t test with Mann-Whitney test was performed. (E) Representative sections showing gene expression in the superior colliculus (SC). *Cnr1*, cannabinoid receptor 1; *Slc32a1*, solute carrier family 32 (GABA vesicular transporter), member 1; *Slc17a6*, solute carrier family 17 (sodium-dependent inorganic phosphate cotransporter), member 6; *CaMKII*α, calcium/calmodulin-dependent protein kinase II alpha; *Dlx5*, distal-less homeobox 5; *Dlx6*, distal-less homeobox 6, displayed from left to right. (F) Neural subtypes cell plots color-coded by gene expression. magenta, *Cnr1*; red, *Slc32a1*; green, *Slc17a6*; cyan, *CaMKII*α; yellow, *Dlx5*; and gray, *Dlx6*. (G) Multiplexed fluorescent *in situ* hybridization of *Cnr1* with *Slc32a1, Slc17a6, CaMKII*α*, Dlx5, Dlx6* (from left to right) in the SC within a single representative section. Scale bar, 500 μm and 50 μm (enlarged view). (H and I) Quantification of neural subtypes (I) and double-positive types, expressed as a percentage of DAPI. Two-Way ANOVA with Sidak’s multiple comparisons post hoc test was performed. Data are presented as mean ± SEM. n=7 male Ctrl group (C), 6 male GCaMP6s group; 7 female Ctrl group, 6 female GCaMP6s group (D); 13 sections from 3 males and 3 females (H, I). ns (not significant), \**p*<0.05, \*\**p*<0.01, \*\*\**p*<0.001, \*\*\*\**p*< 0.0001. For detailed *p* values, refer to Table S1.

Given the heterogeneity of SC neurons^52^, we molecularly characterized the CB_1_R^+^ populations using multiplex fluorescent *in situ* hybridization (FISH) (Table S2). Spatial mapping showed that CB_1_RL neurons predominantly co-expressed glutamatergic markers (VGLUT2 (*Slc17a6*)L: 28.86±4.83%; CaMKIIαL: 26.72±6.62%) but rarely GABAergic markers (VGAT (*Slc32a1*)L: 21.19±3.89%; Dlx5/6L <0.01%) (Figure 1E-I, S2C-D) with no observed sex differences.

### CB_1_R in SC GABAergic neurons governs female-specific defensive homeostasis in risk assessment

To dissociate the role of CB_1_R in SC GABAergic and glutamatergic neurons, CB_1_R-flox mice were bilaterally injected with AAV2/9-Dlx5/6-Cre-GFP (CB_1_R cKO) or AAV2/9-Dlx5/6-GFP (CB_1_R Ctrl) (Figure 2A) and AAV-CaMKLα-Cre-GFP (CaMKLα-cKO) or AAV-CaMKLα-GFP (CaMKLα-Ctrl) (Figure S4A). This enabled selective deletion of CB_1_R in GABAergic (Dlx5/6^+^) or glutamatergic (CaMKLα^+^) neurons (Figure 2B, S4B). Dlx5/6-driven AAVs targeted 94.38±1.94% of GABAergic neurons and 26.52±6.80% of glutamatergic neurons (Figure S5), while CaMKLα-driven AAVs transduced approximately 71±5.33% of glutamatergic neurons (Figure S5D, S5E). Using a hierarchical behavioral analysis framework (HBAF)^51^, we quantified defensive responses across groups (Figure 2C, S4C). In Dlx5/6 CB_1_R cKO mice, most escaped to the refuge after looming stimuli (Figure 2D-2E), while some exhibited alternative strategies like freezing or vigilance (Figure S7, Table S3). Female Dlx5/6 CB_1_R cKO mice exhibited accelerated defensive behavior, with reduced response latency and return time to the refuge (Figure 2E), unlike males (Figure 2D). Refuge duration remained consistent across groups (Figure 2D-E). In contrast, CaMKLα CB_1_R cKO mice exhibited reduced defensive behavior in both sexes, as evidenced by fewer flight-to-refuge events (Figure S4D, E). Males additionally showed shortened refuge time, a phenotype absent in females (Figure S4D, E).

**Figure 2.**
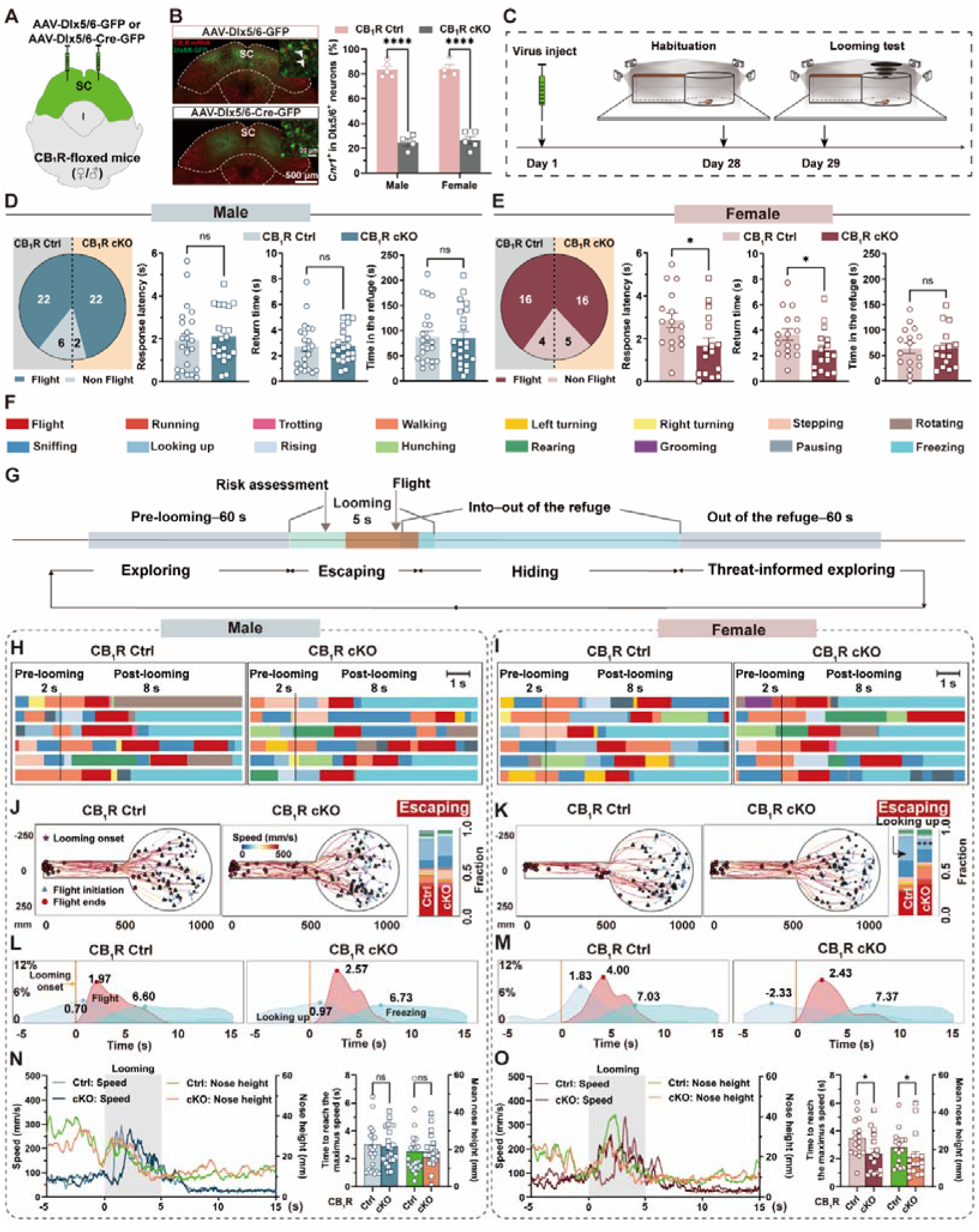
Deletion of CB_1_R in SC Dlx5/6^+^ neurons female-specifically impairs defensive homeostasis in risk assessment. (A) Schematic representation of viral injection of AAV-Dlx5/6-GFP or AAV-Dlx5/6-Cre-GFP into the bilateral SC of CB_1_R-floxed mice. (B) Left: Representative images showing AAV-Dlx5/6-GFP (top) and AAV-Dlx5/6-Cre-GFP (bottom) expression in the SC, alongside CB_1_R mRNA (red) and Dlx5/6^+^ cells bodies (green) in CB_1_R Ctrl and CB_1_R cKO group. Scale bar, 500 μm and 20 μm (enlarged view). White arrows indicate CB_1_R mRNA fluorescent signals colocalized with GFP positive signals. Right: Quantification of CB_1_R^+^ cells in Dlx5/6^+^ neurons in CB_1_R Ctrl and CB_1_R cKO groups for both sexes. (C) Experimental timeline and schematic paradigm of virus injection and the looming test in a refuge-containing apparatus. (D and E) Defensive strategy (left) and flight-to-refuge behaviors (from left to right: response latency, return time, time spent in the refuge) in male (D) and female mice (E) across CB_1_R Ctrl and CB_1_R cKO group. (F) Sixteen movement types during defensive behavior, represented by corresponding colors. (G) Schematic paradigm of the different stages of defensive behavior: exploring, escaping, hiding and threat-informed exploring. (H and I) Ethograms showing the 16 movement types in males (H) and females (I) across groups (CB_1_R Ctrl group on the left, CB_1_R cKO group on the right). Scale bar, 1 s. (J-K) Trajectories (left and middle) and quantification of fractions (right) of mice during the escaping stage (from the onset of looming to the end of flight) in males (J) and females (K) across CB_1_R Ctrl and CB_1_R cKO groups. Symbols indicate mouse locations: five-pointed star for looming onset, triangle for flight initiation, and circle for flight termination. (L and M) Distribution and positions of mice during flight, looking up, and freezing from 5 s pre-looming to 15 s post-looming in males (L) and females (M) across groups (CB_1_R Ctrl on the left, CB_1_R cKO on the right). (N and O) Left: Real-time speed and nose height from 5 s pre-looming to 15 s post-looming in males (N) and females (O) across groups. Right: Quantification of the time taken to reach maximum speed and mean nose height during looming stimuli in males (N) and females (O). Data are presented as mean ± SEM (B, D, E, N, O) or mean (J-O). n=5 male CB_1_R Ctrl, 4 male CB_1_R cKO, 4 female CB_1_R Ctrl, 5 female CB_1_R cKO (B); 28 male CB_1_R Ctrl, 24 male CB_1_R cKO (D), 20 female CB_1_R Ctrl, 21 female CB_1_R cKO (E); 22 male CB_1_R Ctrl and CB_1_R cKO, 16 female CB_1_R Ctrl and CB_1_R cKO (H-O). Two-Way ANOVA with Sidak’s multiple comparisons post hoc test (B, J-K) and Unpaired Student’s t test with Mann-Whitney test (D, E, N, O) were performed. ns (not significant), \**p*<0.05, \*\*\**p*<0.001. For detailed *p* values, refer to Table S1.

HBAF-derived 16-movements classification (Table S4, validated in 3D feature space, Figure 2F, 2H, 2I, S4F) identified four behavioral phases: exploring, escaping, hiding, and threat-informed exploring (Figure 2G, S4G). Movement fractions during pre-looming and hiding in the refuge stage were comparable across groups (Figure S6A-D). Female CB_1_R cKO mice showed impaired risk assessment during escaping, with decreased looking up (Figure 2K), faster flight initiation probability (Figure 2M), as well as reduced nose elevation and time-to-maximum speed (Figure 2O, Movie S1, S2), while males showed no comparable changes (Figure 2J, 2L, 2N). These alterations, including a leftward shift in flight probability distribution (Figure 2M), indicate compromised risk evaluation in female CB_1_R cKO mice. Post-refuge exploration revealed increased sniffing in females (Figure S6F) not in males (Figure S6E), suggesting compensatory environmental sampling. Principal component analysis (PCA) of the 16-dimensional movement space detected no behavioral differences in the four phases (Figure S6G-H). Unlike Dlx5/6 CB_1_R cKO mice, CaMKLα CB_1_R cKO showed no sex-dependent phenotypes during escaping (Figure S4H-4K and Movie S3, S4, Table S5) and hiding (Figure S4L, S4M), likely due to the low specificity (∼70%) of CaMKIIα for glutamatergic neurons. Subsequent experiments focused on SC GABAergic neurons, where CB_1_R exhibits sexually dimorphic regulatory functions.

### CB_1_R^+^ SC-LHb GABAergic circuit governs female-specific defensive homeostasis in risk assessment

To identify the specific circuits through which downstream SC CB_1_R^+^Dlx5/6^+^ neurons mediate female risk assessment during defensive behavior, we conducted anterograde neuronal tracing from CB_1_R^+^Dlx5/6^+^ neurons in the SC. This revealed over 30 projection targets without sex differences in fiber density (Figure S8). Parallel cell-specific labeling showed comparable CB_1_R^+^ Dlx5/6^+^ neurons density between sexes (Figure S9), highlighting conserved circuit architecture. c-Fos screening revealed that LHb neurons were selectively activated by looming stimuli in females following SC CB_1_R ablation in Dlx5/6 neurons (Figure 3A-3D). Given the LHb’s established role in aversive processing^53–60^ and eCB-mediated coping^61^, we hypothesized that the CB_1_R^+^ SC-LHb GABAergic circuit regulates female risk assessment. Retrograde tracing confirmed dense SC CB_1_R^+^ Dlx5/6^+^ axons innervation of the LHb (Figure S10). Fiber photometry recordings used to detect eCB dynamics via the eCB sensor^62–64^ (Figure 3E) showed looming stimuli increased eCB and calcium signals in the unilateral LHb of flight but not non-flight-responding mice (Figure 3F-3I), with no sex differences in signal dynamics (Figure 3J). These results suggest that looming induces elevated post-synaptic neuronal activity and eCB release in downstream nuclei in both sexes, specifically in the active flight response. LHb-specific CB_1_R antagonism female-specifically impaired risk assessment (Figure 3K-3N), unlike interventions in reuniens thalamic nucleus or zona incerta (ZI) (Figure S11).

**Figure 3.**
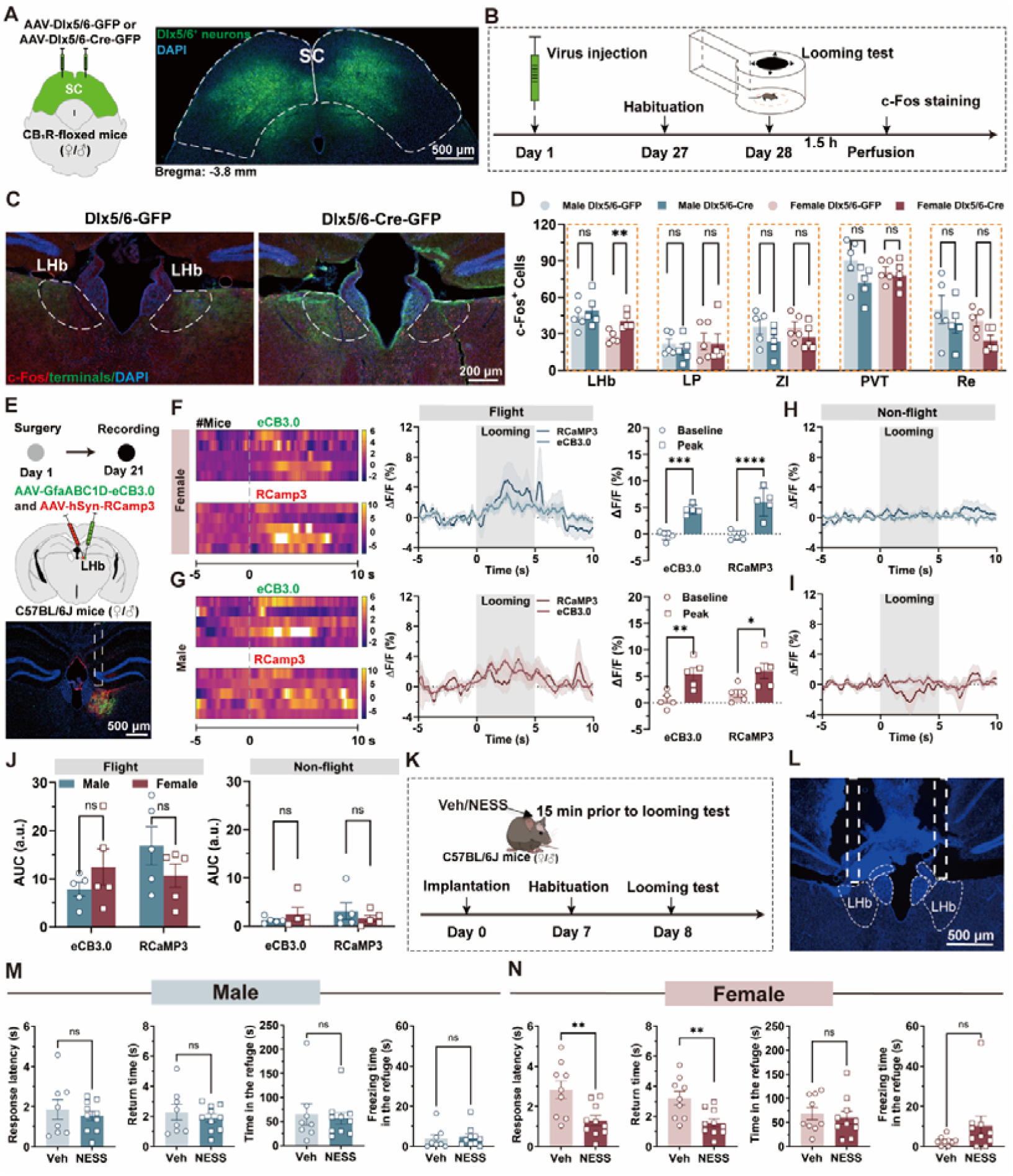
LHb serves as a key downstream target of the SC that mediates female-specific modulation of defensive homeostasis in risk assessment. (A) Left: Schematic representation of viral injections of AAV-Dlx5/6-GFP or AAV-Dlx5/6-Cre-GFP into the bilateral SC of CB_1_R-floxed mice. Right: Representative images showing Dlx5/6^+^ GFP signals expression in the SC. Scale bar, 500 μm. (B) Experimental timeline and schematic paradigm of the looming test. (C) Representative images of c-Fos expression in the LHb in response to looming stimuli in female CB_1_R Ctrl and CB_1_R cKO mice. Scale bar, 200 μm. (D) Quantification of c-Fos expression in different brain regions across sexes. Abbreviations: LHb, lateral habenula; LP, lateral posterior thalamus; ZI, zona incerta, PVT, paraventricular thalamic nucleus; Re, reuniens thalamic nucleus. (E) Experimental timeline (top), schematic paradigm (middle) and representative image (bottom) of the viral infection in LHb neurons. Scale bar, 500 μm (F, G) Heatmap visualization of endocannabinoid (eCB) and calcium (lower panel) dynamics (left), signals (middle), and peak signals (right) in male (F) and female (G) mice during flight response to looming stimuli. Shaded areas around the means indicate error bars; the gray shadow marks the looming stimuli event. (H, I) eCB and calcium fluorescence changes in male (F) and female (G) mice during non-flight response to looming stimuli. (J) Statistical graph showing the area under the curve (AUC) of eCB and calcium signaling in male and female mice with flight (left) or non-flight (right) response under looming stimulus. (K) Timeline and schematic representation of pharmacological experiment during the looming test. (L) Representative image showing the placement of track cannula in the bilateral LHb. Scale bar, 500 μm. (M and N) Metrics of defensive behavior (from left to right: response latency, return time, time spend in the refuge and freezing time in the refuge after looming stimuli) for male (M) and female mice (N) in vehicle (Veh) and NESS 0327 (NESS) groups. Data are presented as mean ± SEM. n=5 per group (D), 4 per group (H), 5 per group (F-J), 8 male Veh, 10 male NESS (K), 9 female Veh, 10 female NESS (L). Unpaired Student’s t test with Mann-Whitney test (D, K, L) and two-way ANOVA with Sidak’s multiple comparisons post hoc test (H) were performed. ns (not significant), \**p*<0.05, \*\**p*<0.01, \*\*\**p*<0.001, \*\*\*\**p*<0.0001. For detailed *p* values, refer to Table S1.

Optogenetic activation of CB_1_R^+^ SC-LHb GABAergic projections (Figure 4A-4D) disrupted female risk assessment (Figure 4F) but not males (Figure 4E), while chemogenetic inhibition (Figure 4G-4I) prolonged female risk assessment (Figure 4K), evidenced by increased latency to initiate flight and return to the refuge. These effects were not observed in CB_1_R^+^ GABAergic SC-ZI (Figure S12A-S12D) or SC-lateral posterior thalamus (LP) (Figure S12E-S12H) protections, confirming the circuit’s necessity. Male defensive behavior (Figure 4J) and spontaneous behavior in both sexes remained unaffected (Figure 4L-P, Table S6). In summary, CB_1_R in SC GABAergic projections to the LHb are necessary for female to exhibit full, deliberative risk assessment comparable to males. Their genetic deletion compromised defensive homeostasis in risk assessment, manifesting shortened and inadequate threat evaluation prior to flight.

**Figure 4.**
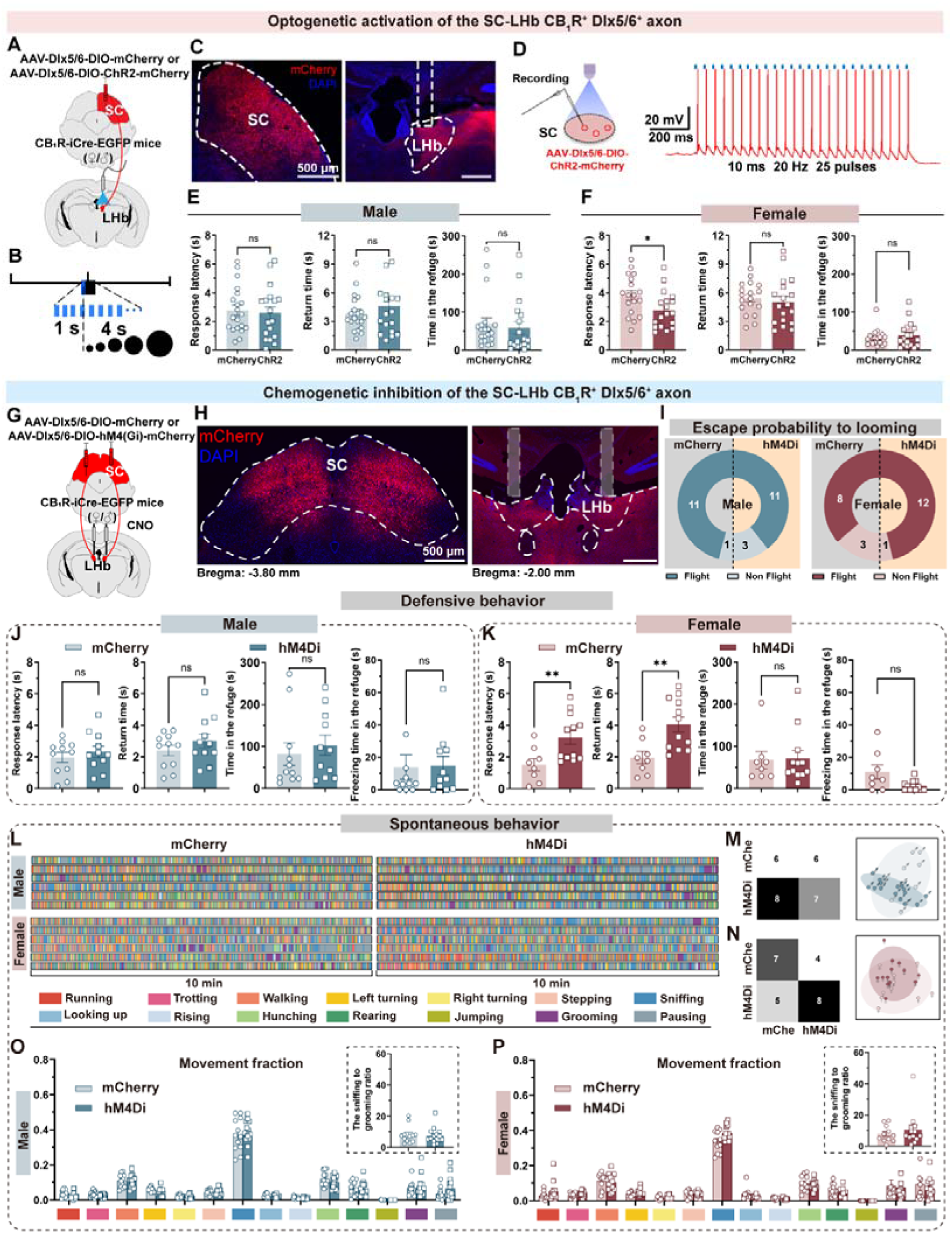
Modulating SC-LHb CB_1_R^+^Dlx5/6^+^ projections female-specifically affects defensive homeostasis in risk assessment. (A) Schematic representation of unilateral blue light activation targeting CB_1_R**^+^**SC-LHb Dlx5/6^+^ projections. (B) Protocol for optogenetic stimulation. (C) Representative image showing ChR2 virus expression in the SC of a CB_1_R-iCre-EGFP mouse and the location of the fiber track (blue, DAPI; red, ChR2-mCherry; dotted border line, fiber track). Scale bars, 500 μm. (D) Left: Schematics of patch-clamp slice recording during optogenetic stimulation of SC CB_1_R^+^Dlx5/6^+^ neurons. Right: Light-pulse train triggered spikes (red) in ChR2-mCherry–positive neurons. (E and F) Metrics of defensive behavior in males (E) and females (F) (from left to right: response latency, return time and time spend in the refuge after looming) under optogenetic activation of CB_1_R**^+^** SC-LHb Dlx5/6^+^ terminals. (G) Schematic representation of bilateral chemogenetic inhibition targeting CB_1_R**^+^**SC-LHb Dlx5/6^+^ projections. (H) Representative image showing CB_1_R^+^Dlx5/6^+^ neurons in the SC and the fiber track location (blue, DAPI; red, ChR2-mCherry; dotted border line, fiber track). Scale bars, 500 μm. (I) Defensive strategy in males (left) and females (right). (J and K) Metrics of defensive behavior (from left to right: response latency, return time, time spent in the refuge, and freezing time in the refuge after looming stimuli) in male (J) and female mice (K) in mCherry and hM4Di group. (L) Ethograms depicting 14 movements in males (top) and females (bottom) across different groups (mCherry and hM4Di group) in spontaneous behavior. Each color corresponds to a specific movement. (M, N) A random forest classifier trained on 14-dimensional movement fractions to predict group labels (left), and PCA visualization of behavioral features (right) for male (m) and female (n) in the mCherry and hM4Di groups in spontaneous behavior. (O, P) Comparison of the movement fraction, the sniffing to grooming ratio between male (L) and female (M) mice in the mCherry and hM4Di groups. Data are presented as mean ± SEM. n=20 male mCherry, 18 male ChR2 (E), 18 female mCherry and ChR2 (F),11 male mCherry and hM4Di (J), 9 female mCherry, 12 female hM4Di (K). n=12 male mCherry group, 14 male hM4Di group (O); 11 female mCherry group, 13 female hM4Di group (P). Two-Way ANOVA with Sidak’s multiple comparisons post hoc test was performed (O, P). Unpaired Student’s t test with Mann-Whitney test was performed. ns (not significant), \**p*<0.05, \*\**p*<0.01. For detailed *p* values, refer to Table S1.

### CB_1_R-mediated suppression of GABA release from SC terminal onto the LHb underlies female-specific phenotypes

Given that presynaptic CB_1_R inhibits neurotransmitter release^65,66^, we measured GABA dynamics in the LHb using a genetically encoded fluorescent sensor for *in vivo* imaging (Figure 5A-5C). Female Dlx5/6 CB_1_R cKO mice exhibited elevated GABA release during looming (Figure 5E), unlike males (Figure 5D), indicating disinhibited transmission from CB_1_R^+^ SC GABAergic projections to the LHb. The LHb expressed high levels of diacylglycerol lipase alpha (Dagla) (Figure 5G), essential for synthesizing 2-arachidonoylglycerol (2-AG)^67,68^. Locally deletion of *Dagla* by bilaterally injecting AAV-EF1α-Cre-EGFP virus into the LHb of Daglα-floxed mice (Figure 5F-5I) induced female-specific risk assessment deficits (Figure 5K), similar to those seen in Dlx5/6 CB_1_R cKO mice, with no equivalent effects in males (Figure 5J). These findings suggest that 2-AG retrograde signaling in the LHb gates presynaptic GABA release from CB_1_R^+^ SC GABAergic terminals, thereby ensuring that females maintain defensive homeostasis in risk assessment comparable to males. Disruption of this circuit through CB_1_R knockout, circuit modulation, or 2-AG depletion consistently impairs female-specific defensive homeostasis in threat evaluation.

**Figure 5.**
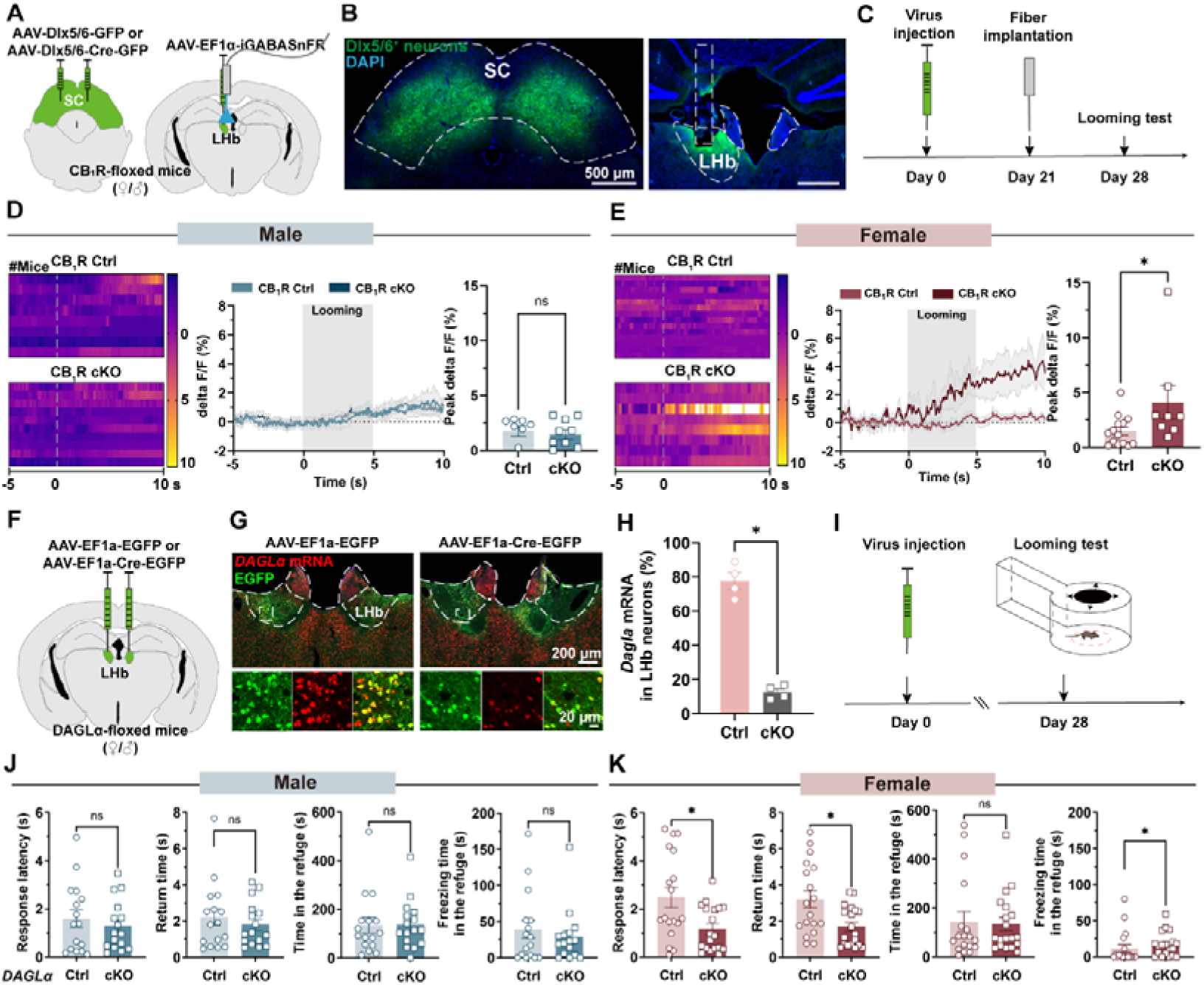
SC-LHb GABA transmission is required for risk assessment in a female-specific manner. (A) Schematic representation of viral injection: AAV-Dlx5/6-GFP or AAV-Dlx5/6-Cre-GFP into the bilateral SC (left) and AAV-EF1α-iGABASnFR into the unilateral LHb (right) of CB_1_R-floxed mice. (B) Representative image showing Dlx5/6^+^ GFP signals in the SC and iGABASnFR signals in the LHb, and the position of the fiber track (blue, DAPI; green, Dlx5/6^+^ GFP or iGABASnFR signals; dotted border line, fiber track). Scale bars, 500 μm. (C) Apparatus for looming test and schematic of the recording system to measure GABA signals using fiber photometry during looming stimuli. (D and E) Heatmap of iGABASnFR fluorescence changes (left), signals (middle), and peak signals (right) in male (D) and female (E) mice. Shaded areas around the means indicate error bars; the gray shadow marks the looming stimuli event. (F) Schematic representation of viral injection: AAV-EF1α-EGFP or AAV-EF1α-Cre-EGFP into the unilateral LHb (left) of Daglα-floxed mice. (G) Representative images showing AAV-EF1α-EGFP (left) and AAV-EF1α-Cre-EGFP (right) expression in the LHb (red, Daglα mRNA; green, EGFP signals). Scale bar, 200 μm and 20 μm (enlarged view). (H) Percentage of DAGLα^+^ cells in LHb neurons in DAGLα Ctrl and DAGLα cKO group. (I) Schematic paradigm for the looming test. (J and K) Metrics of defensive behavior (from left to right: response latency, return time, time spent in the refuge and freezing time in the refuge after looming stimuli) of male (J) and female mice (K) in DAGLα Ctrl and DAGLα cKO groups. Data are presented as mean ± SEM. n = 8 male CB_1_R Ctrl, 9 male CB_1_R cKO (D), 14 female CB_1_R Ctrl, 8 female CB_1_R cKO (E), 4 per group (H), 16 male DAGLα Ctrl and DAGLα cKO (J), 18 female DAGLα Ctrl, 19 female DAGLα cKO (K). Unpaired Student’s t test with Mann-Whitney test. ns (not significant), \**p*<0.05. For detailed *p*-values, refer to Table S1.

### The loss of CB_1_R in SC GABAergic neurons underlies a female-specific vulnerability to stress

Considering that anxiety can accelerate defensive responses to looming stimuli^69^, we assessed rat stress susceptibility in SC GABAergic CB_1_R cKO and Ctrl mice (Figure 6A). Chronic stress resulted in comparable weight stability across groups (Figure 6B), but female Dlx5/6 CB_1_R cKO mice showed increased anxiety-like behavior, evidenced by reduced open-arms time in elevated plus maze post-stress (Figure 6C). Stress-induced behavioral reorganization included reduced locomotion (Figure 6G, 6H, 6J, 6K, Table S6), increased vertical exploration, such as rearing, with decreased nose height (Figure 6I, 6L), and decreased horizontal exploration, such as sniffing, with increased nose height (Figure 6I, 6L) in male and female CB_1_R Ctrl and cKO groups. The stressor elicits a strategic behavioral shift from active exploration to defensive vigilance, manifesting as reduced locomotion and heightened vertical investigation, which potentiates visual information acquisition for environmental surveillance. Female CB_1_R cKO mice exhibited a pathological triad post-stress, characterized by anhedonia-like reduced sniffing (10**–**20 min) (Figure 6L), hypervigilance associated with increased corner occupancy (40**–**50 min) (Figure 6K), and compulsion-like elevated grooming (50**–**60 min) (Figure 6L). This progression resembles human anxiety disorders (DSM-5), confirming female-specific vulnerability. Machine learning analysis quantified significant behavioral dissimilarity induced by stress (Figure S13, S14). Collectively, CB_1_R deletion in SC Dlx5/6^+^ neurons triggers a female-specific pathogenic cascade characterized by elevated GABA release in the LHb, impaired risk assessment, and heightened vulnerability to stress-induced psychopathology. This defines an evolutionarily conserved circuit that governs sex-specific survival plasticity through a vertebrate-conserved neuromodulatory system.

**Figure 6.**
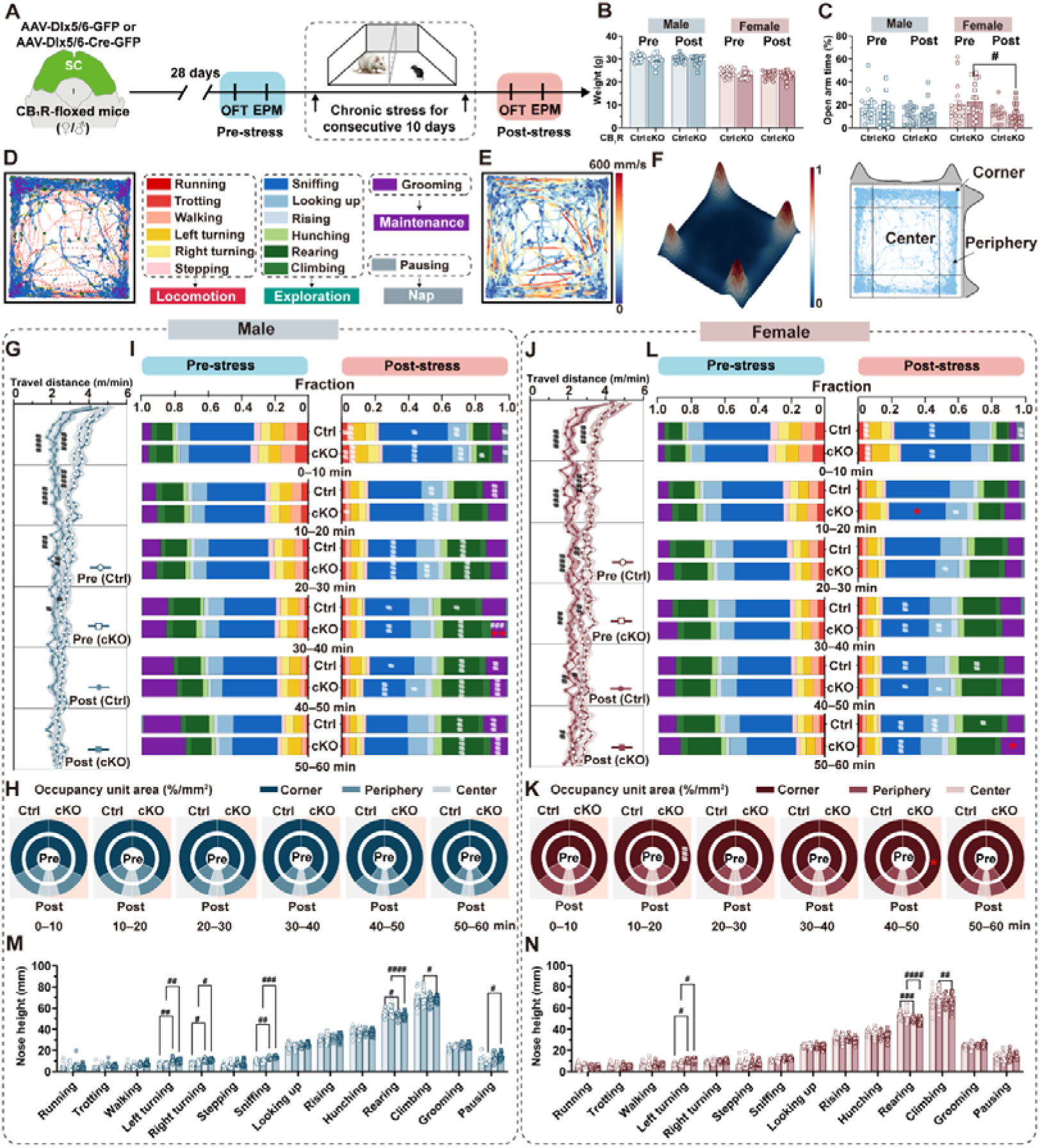
CB_1_R deletion in SC GABAergic neurons increases abnormal behavior patterns specifically in females after stress. (A) Schematic representation of viral injection of AAV-Dlx5/6-GFP or AAV-Dlx5/6-Cre-GFP into the bilateral SC of CB_1_R-floxed mice and the timeline of behavioral teats conducted pre- and post-chronic stress. (B, C) Changes in body weight and percentage of time spent in open arms during elevated plus maze test of mice pre- and post-chronic stress across different groups. #*p*<0.05. (D) Trajectories showing speed during 60-min of open field activity. (E) Representative trajectories illustrating different movements in the open field apparatus. Each of the 14 colors corresponds to a specific movement type. (F) Density map showing the spatial distribution of dorsal points during 60-min of open field (left) and representative trajectories in the center, periphery, and corner (right) of the apparatus. (G, J) Travel distance per minute throughout the experiment for males (G) and females (J). (I, L) Comparison of the fraction (10-min epochs) between male (I) and female (L) mice in the CB_1_R Ctrl and CB_1_R cKO groups. (H, K) Pie charts showing relative percentage of time spent in different areas for males (H) and females (K). Inner and outer rings represent pre- and post-stress percentages, respectively, while left and right halves correspond to CB_1_R Ctrl (left) and CB_1_R cKO (right) groups. (M, N) Nose height during different movements in the 60-min open field apparatus (bottom) for males (M) and females (N). Data are presented as mean ± SEM (B, C, G, J, M, N) and as mean (I, L, H, K). n=18 pre-stress, 17 post-stress male CB_1_R Ctrl; 18 pre-stress, 16 post-stress male CB_1_R cKO; 17 pre-stress, 20 post-stress female CB_1_R Ctrl; 17 pre-stress, 18 post-stress female CB_1_R cKO. Two-Way ANOVA with Sidak’s multiple comparisons post hoc test was performed. #*p*<0.05; ##*p*<0.01; ###*p*<0.001; ####*p*<0.001 pre-stress versus post-stress; \**p*<0.05; \*\**p*<0.01 post-stress CB_1_R Ctrl versus CB_1_R cKO For detailed *p* values, refer to Table S1.

## DISCUSSION

This study identifies a female-specific CB_1_R-gated SC-LHb GABAergic circuit that orchestrates defensive homeostasis in risk assessment. We demonstrate how evolutionarily conserved eCB signaling enables subsecond-scale precision in risk assessment through sexually dimorphic modulation of a subcortical pathway. Our findings establish a neural mechanism whereby life-history trade-offs are implemented at the circuit level, prioritizing female survival through specialized neuromodulatory control of threat processing.

### CB_1_R as a sexually dimorphic neuromodulator in the homeostatic maintenance of risk assessment

CB_1_R, a key component of the evolutionarily conserved eCB system across vertebrates^40^, modulates various innate behaviors^70,71^ and exhibits sexual differences in neural function^72^. Although associated with locomotor activity^63,73,74^, nociceptive processing^75^ and affective states^45,76^, here we reveal that CB_1_R in the SC are necessary for visual threat-induced defensive behavior in both sexes, expanding previous studies upon its role in defensive behavior^77,78^. Notably, although visual threats activate SC CB_1_R^+^ neurons universally, CB_1_R-mediated modulation of the SC-LHb GABAergic circuit specifically governs risk computation in females. This dissociation between conserved neural activation and sex-specific neuromodulation^79^ reveals a previously unrecognized organizational principle for implementing sexually dimorphic behaviors.

The functional compartmentalization of eCB signaling observed here extends previous demonstrations of nucleus-^80–82^ and cell type-specific^83–88^. Specifically, a recent study revealed that 2-AG acts on neurons to enhance synaptic inhibition, while anandamide targets astrocytes to promote synaptic potentiation^89^. This cell type- and endocannabinoid-specific signaling paradigm provides critical support that eCB signaling achieves functional specialization, including the sex-specific modulation of the SC-LHb GABAergic circuit observed here, by leveraging distinct molecular and cellular targets. Despite conserved anatomical projections between the SC and LHb, CB_1_R affect GABAergic transmission within this pathway exclusively in females, facilitating more deliberate threat evaluation, an adaptation likely driven by evolutionary pressures that prioritize female survival. This specialization contrasts with its non-dimorphic role in glutamatergic circuits regulating escape behavior^34,90–93^, whereas GABAergic neurons provide inhibitory control over these responses^94^. Such functional segregation may reflect evolutionary optimization where conserved subcortical pathways are overlaid with sex-specific modulatory controls to accommodate different survival strategies.

GABAergic neurons orchestrate behaviors by regulating excitation-inhibition balance^95^, gating signal throughput^96^, and facilitating disinhibition^97^, which allows for precise control over the intensity, direction, and timing. Different GABAergic subtypes across brain regions encode specific fear behavior^98^, regulate defensive outputs^90^, and coordinate vigilance^99,100^. Critically, these GABAergic functions themselves are often sexually dimorphic^101^, contributing to sex differences in defensive behavior. Thus, these findings suggest that CB_1_R modulates defensive behavior through complementary mechanisms, orchestrating behavioral initiation via glutamatergic regulation in both sexes, while maintaining defensive homeostasis through sexually dimorphic modulation via GABAergic circuits. This dual arrangement may support immediate individual survival and long-term species-level reproductive strategies.

This functional dimorphism mediated via CB_1_R parallels mechanisms such as tachykinin receptor 1 activation in preoptic area of the hypothalamus neurons, which drives male-specific mating behavior^19^. Both findings highlight how evolutionarily distinct neuromodulatory systems, the endocannabinoid and substance P systems, can be deployed within specialized circuits to promote sex-specific behaviors. We propose that such dimorphism may originate from early developmental programming, as eCB signaling has been shown to regulate microglia-dependent synaptic pruning and astrogenesis, thereby shaping sex differences in juvenile social behavior^102^. Further supporting a developmental basis, endocrine signals have been shown to influence the connectivity and gene expression profiles of hormone-responsive neurons^31^, creating latent sex biases that are selectively activated by specific threat contexts in adulthood.

### Subcortical GABAergic circuit architecture governs sex-specific threats evaluation

Subcortical circuit governing sex-specific survival strategies exhibit conserved anatomy alongside different neuromodulation. While sexually dimorphic neural architectures are well-documented in hypothalamic^20,103^ and limbic hubs^104^, manifesting as sex differences in gene expression^105^, synaptic activity^106,107^, and cell density^103^, the SC-LHb circuit demonstrates functional sexual dimorphism without structural differences, as previously reported^52^ and confirmed here. The SC, a conserved detector of survival-relevant visual threats across vertebrates, processes threat information through glutamatergic neurons^34,90–93,108^ and GABAergic neurons^94^. Our findings reveal a specific sexually dimorphic mechanism within the CB_1_R^+^ SC-LHb GABAergic circuit. We show that female-specific activation of the LHb depends on CB_1_R-mediated retrograde 2-AG signaling that selectively enhances presynaptic inhibition of GABA release. This neuromodulatory specialization aligns with life-history trade-offs that prioritize female survival due to greater reproductive investment.

The LHb is broad involvement in encoding aversion, anxiety, and stress^54,109,110^, positions it as a strategic hub for integrating female-specific survival behaviors. Our work identifies that the LHb integrates female-specific survival behaviors. Its broad involvement across these behaviors suggests that the LHb may function as a central integrator of survival-stress information, with sex differences in connectivity^53,111^ potentially underlying differential susceptibility to stress-related disorders. This integrative function may also account for the heightened threat reactivity often seen in individuals with emotional disorders. Beyond risk assessment, the LHb also contributes to risk preference evaluation^112^ and parental behavior^113,114^ via distinct afferent pathways, highlighting its broader function in balancing survival-reproduction trade-offs. The engagement of these functions in the LHb highlights its central role in female-specific adaptive behaviors, with eCB signaling providing essential neuromodulatory flexibility to dynamically prioritize competing demands, based on internal and external contexts.

More broadly, our results align with growing evidence that subcortical structures, including the BNST^19,24,115^ and VMH^18,21,31^, encode sexually dimorphic behaviors in a cell-type-specific mechanisms involving neuronal populations expressing Tac1^19^, serotonin^115^, aromatase^24^, or progesterone-receptor^18,21^. This supports a unifying paradigm in which evolutionarily conserved subcortical circuits use genetically specified neuronal subpopulations to regulate innate behaviors in a sex-dependent manner. Such functional specialization, often mediated by inhibitory signaling, is conserved across species^116–119^ while allowing for species-specific adaptations. Importantly, dysregulation of this circuits can disrupt defensive homeostasis, as evidenced by the LHb’s involvement in female-biased disorders such as PTSD^120,121^ and postpartum anxiety^122,123^, highlighting the clinical relevance of these evolutionarily tuned circuits.

### Defensive homeostasis, evolutionary trade-offs and therapeutic translation for sex-biased disorders

The behavioral homeostasis theory suggests that the evolutionary function of defensive behavior is to rapidly maximize an organism’s overall readiness to cope with new survival threats and to minimize unnecessary energy expenditure^124–126^. This concept aligns with recently identified homeostatic systems regulating fundamental needs such as sleep^127^ and social interaction^128^. Building on our previous findings that complete defensive behavior includes not only threat detection and escape to shelter, but also the critical process of re-emergence for risk reassessment and environmental exploration^30^, we introduce the concept of defensive homeostasis. This homeostatic process involves the entire sequence of responses to life-threatening stimuli, emphasizing the optimization of an organism’s ability to rapidly evaluate whether external stimuli present imminent survival risks. Notably, such homeostatic systems often exhibit inherent sexual dimorphism, likely rooted in genetic and epigenetic mechanisms—as seen in the sex-specific imprinting of hypothalamic genes^129^ which may predispose differential defensive strategies between males and females. Although both sexes showed consistent escape responses to immediate threats, females displayed quicker re-exploration and more thorough risk reassessment after reaching safety^30^. This enhanced defensive homeostasis in females can be understood through the life-history trade-off theory proposed by William Williams in 1957. Due to asymmetric parental investment, with females bearing the costs of gestation and lactation, female behavioral strategies prioritize survival through more cautious and comprehensive risk assessment. We propose that the comparable assessment duration, between sexes served in females as a proactive phase for gathering and processing more detailed information, thereby \ qualitatively enhancing their defensive homeostasis. The reduction of this duration following CB_1_R deletion in SC GABAergic neurons highlights the role of CB_1_R in facilitating the deliberative processing characteristic of female defensive strategy.

Chronic stress exposure further highlights these evolutionary trade-offs, where CB_1_R dysfunction transforms adaptive vigilance into female-specific psychopathology. Consistent with the role of CB_1_R in managing acute^130^ and chronic stress^45,131^, Dlx5/6 CB_1_R-KO mice exhibit maladaptive behaviors exclusively in females, including reduced exploration, heightened thigmotaxis, and compulsive grooming. These phenotypes closely align with the core symptoms of anxiety and depression as outlined in the DSM-5. This maladaptation may arise from disrupted threat calibration within the CB_1_R^+^ SC-LHb GABAergic circuit. From an evolutionary perspective, this vulnerability reflects the costs of optimizing female survival strategies. Females with CB_1_R dysfunction may overestimate risks, a strategy that could theoretically enhance offspring protection during maternal investment^132^ but imposing significant energetic costs. Conversely, males maintain more accurate defensive homeostasis in risk assessment, supporting competitive resource acquisition and reproductive success^13^. These sexually dimorphic behavioral strategies collectively enhance species survival and reproductive adaptability. Clinically, aberrant risk calibration is mechanistically linked to disorders predominantly affecting females, such as pathological hypervigilance in post-traumatic stress disorder (PTSD)^133^, which correlated with habenular hyperactivity^134,135^. Thus, enhancing CB_1_R signaling in this circuit presents a promising therapeutic approach for addressing sex-specific psychopathologies.

### Limitations of the study

The Dlx5/6 promotor exhibits approximately 20% glutamatergic specificity, yet fails to fully resolve the heterogeneity among GABAergic subpopulations in the SC. Future studies should employ intersectional genetics, such as Cre/Flp-dependent transgenic mice with viral tools, for subpopulation-specific manipulation. Additionally, the continuous stress modeling protocol limited the dissection of stage-specific adaptations. Future designs should incorporate *in vivo* recordings during chronic stress exposure to characterize temporal dynamics. Finally, therapeutic translation warrants testing CB_1_R-positive allosteric modulators^136^ for rescuing female-specific deficits.

## STAR ★ METHODS

Detailed methods are provided in the online version of this paper and include the following:

- KEY RESOURCES TABLE
- **RESOURCE AVAILABILITY**

- Lead contact
- Materials availability
- Data and code availability
- EXPERIMENTAL MODEL AND PARTICIPANT DETAILS

- Animals
- Viruses
- METHOD DETAILS

- Experimental design and randomization
- Stereotaxic surgeries
- Cannula implantation
- Drugs delivery
- Histology
- Behavioral assays
- 3D Behavioral data processing
- Chronic rat stress model
- Patch-clamp electrophysiology
- Optical fiber implantation
- Optogenetics
- Fiber photometry
- QUANTIFICATION AND STATISTICAL ANALYSIS

- Statistics

## ACKNOWLEDGMENTS

This work was supported by the Brain Science and Brain-like Intelligence Technology-National Science and Technology Major Project (2022ZD0208300 to F.W.), the National Natural Science Foundation of China (32371062 to F.W., 31630031 and 31930047 to L.W, Research Fund for International Senior Scientists T2250710685), the Guangdong Provincial Key Laboratory of Brain Connectome and Behavior (2023B1212060055 to L.W.), the Guangdong Basic and Applied Basic Research Foundation (2022A1515110120 to X.L.), the Shenzhen Key Basic Research Project (JCYJ20220818100805013 to F.W.), and the Financial Support for Outstanding Talents Training Fund in Shenzhen (to L.W.). Behavioral experiments focusing on chronic stress was conducted at the Shenzhen Brain Science Infrastructure. We are grateful to Dr. Xuemei Liu for conducting the preliminary experiments. We would like to express our sincere gratitude to the members of Feng Wang’s lab for their helpful comments and kind suggestions regarding the manuscript.

## AUTHOR CONTRIBUTIONS

F.W. and L.W. conceived the project, designed the experiments and proposed the concept. X.Liu and H.H. conducted the majority of experiments and analyzed the corresponding data. X.F. and X.Li conducted the optogenetic, morphological, and pharmacological experiments and analyzed the corresponding data. X.Liu and Z.G. conducted the rat stress experiments. J.Y., H.H., X.Liu, and Z.L. contributed to 3D behavioral analysis. R.C., S.C., and Y.L. developed and provided eCB3.0 sensor. H.Y. conducted the RCamp3 and eCB3.0 sensor recordings under the supervision of Z.W.. F.W., X.Liu, H.H., and L.W. wrote the manuscript with input from all authors. L.W. and F.W. supported all aspects of this project.

## DECLARATION OF GENERATIVE AI AND AI-ASSISTED TECHNOLOGIES IN THE WRITING PROCESS

During the preparation of this work, the AI tool DeepSeek and Monica were used to assist with language editing. Upon its use, the authors thoroughly reviewed and further edited the entire text. The authors take full responsibility for the content of the publication.

## STAR ⍰ METHODS

Detailed methods are provided in the online version of this paper and include the following:

## KEY RESOURCES TABLE

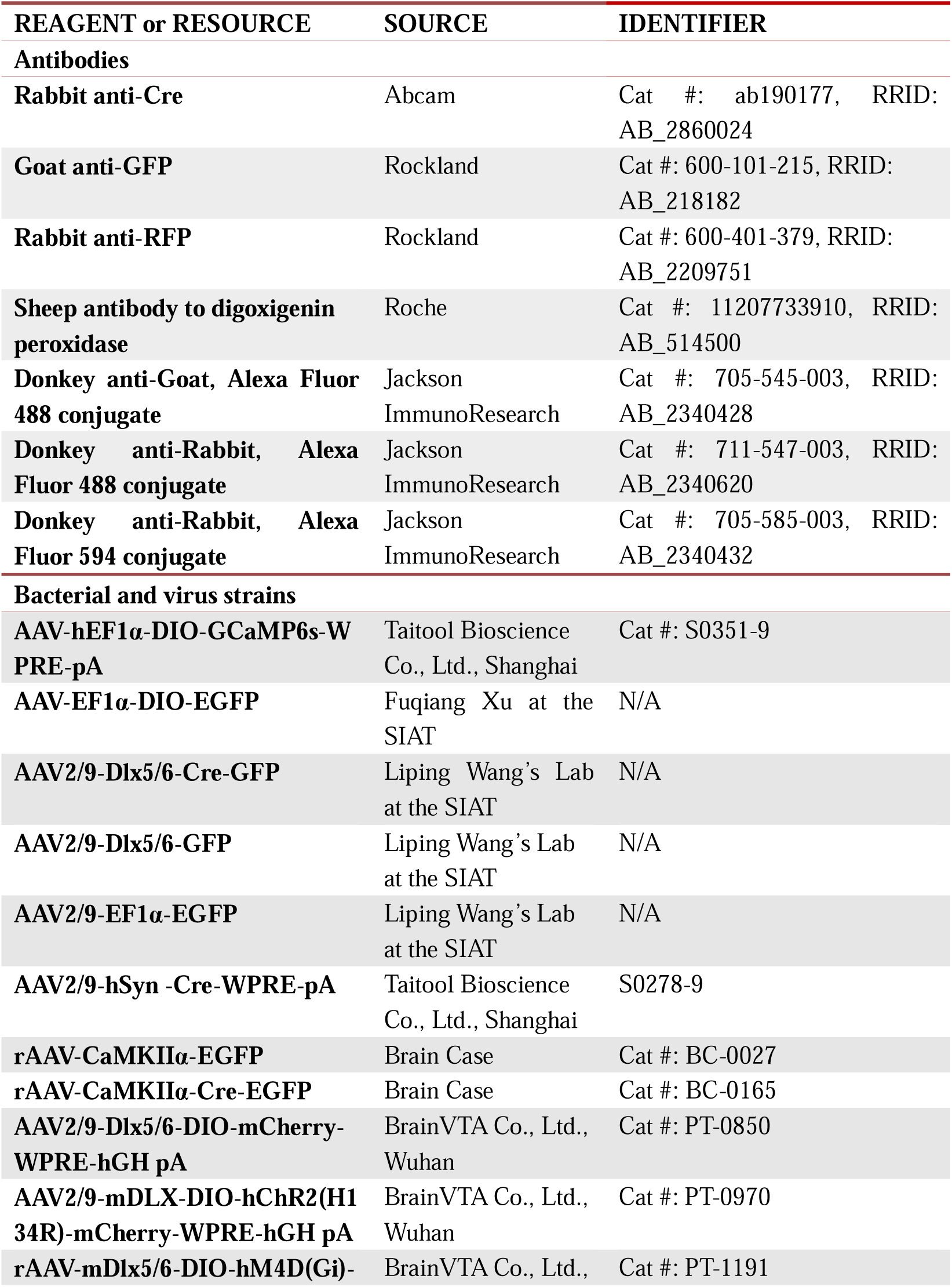
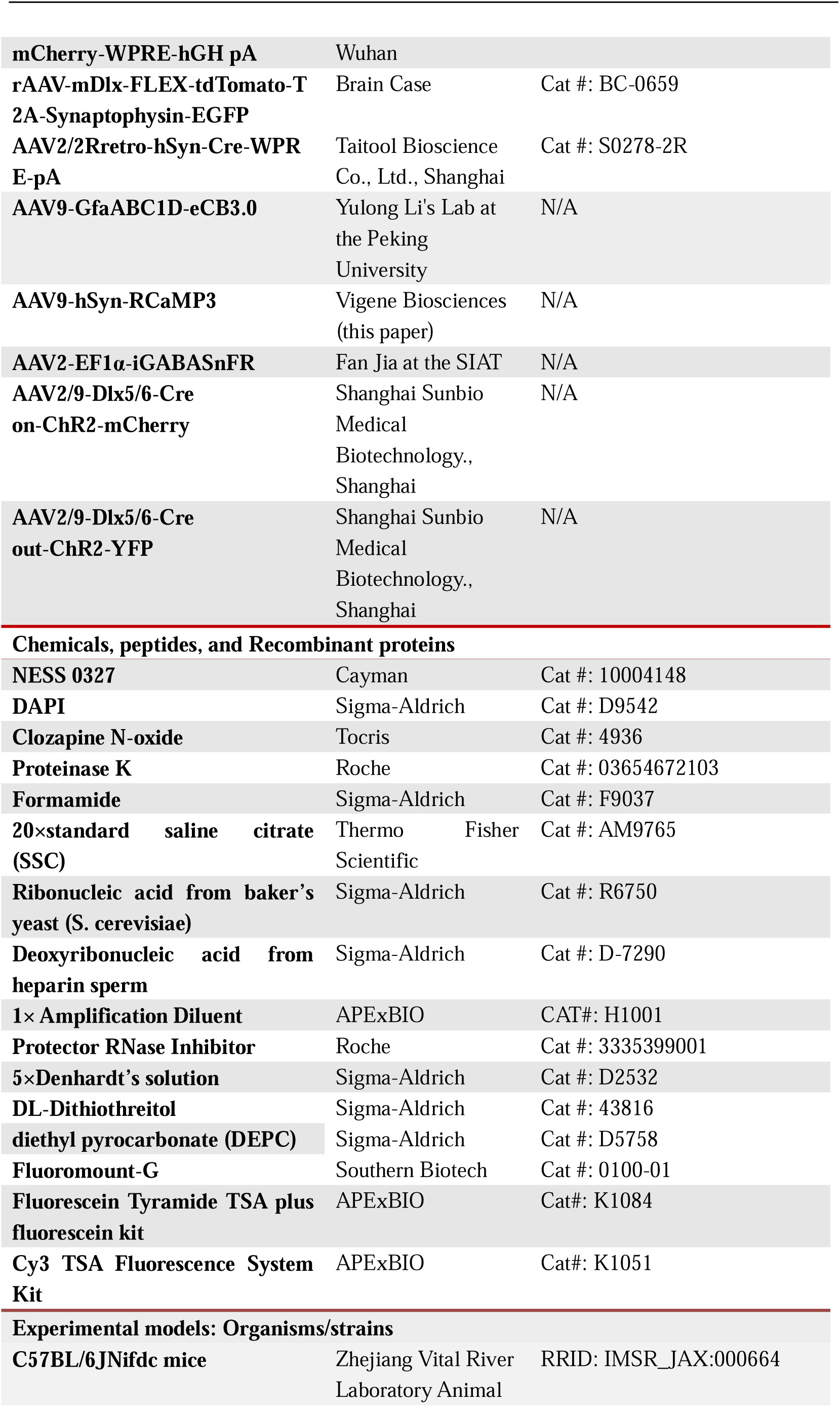
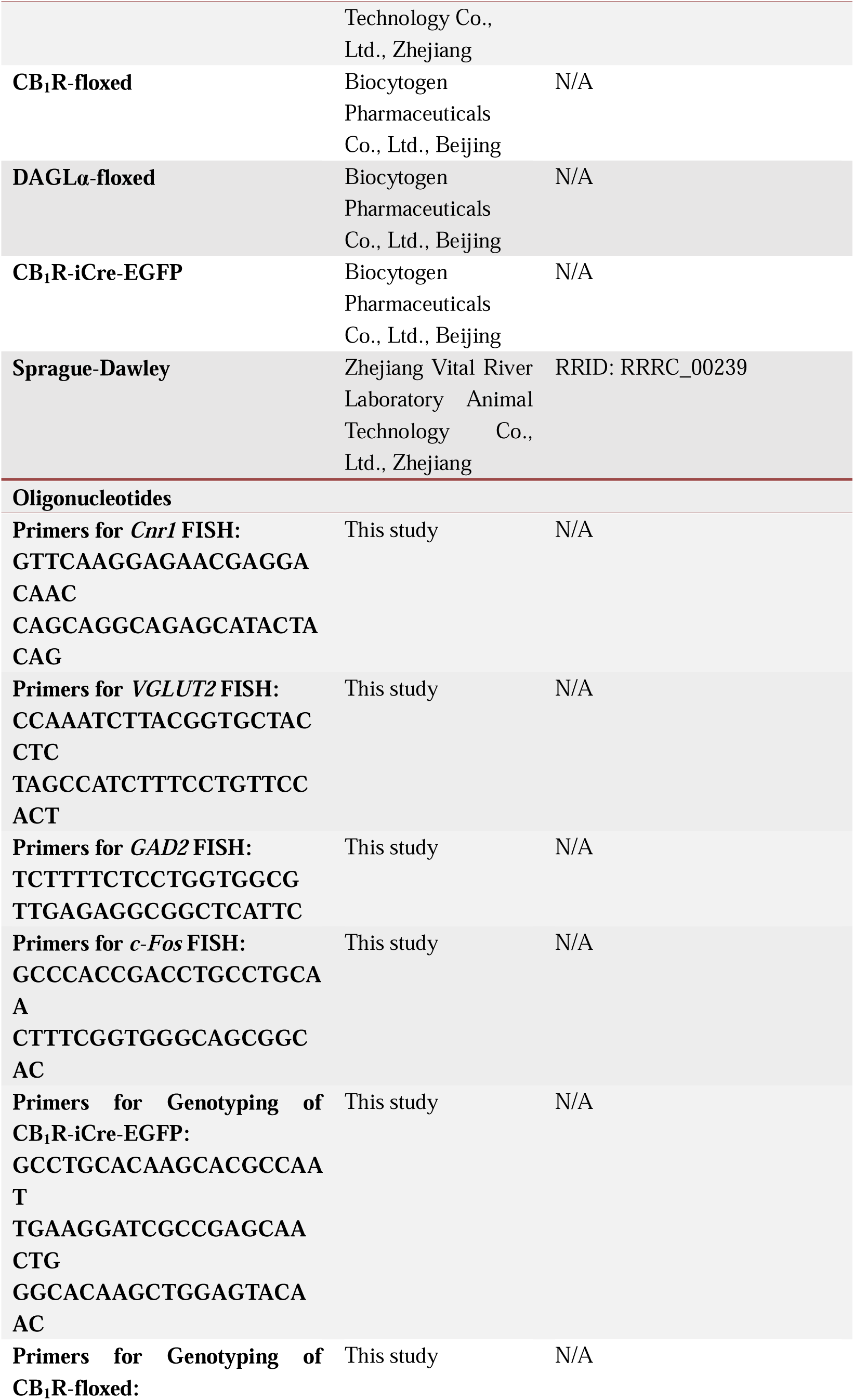
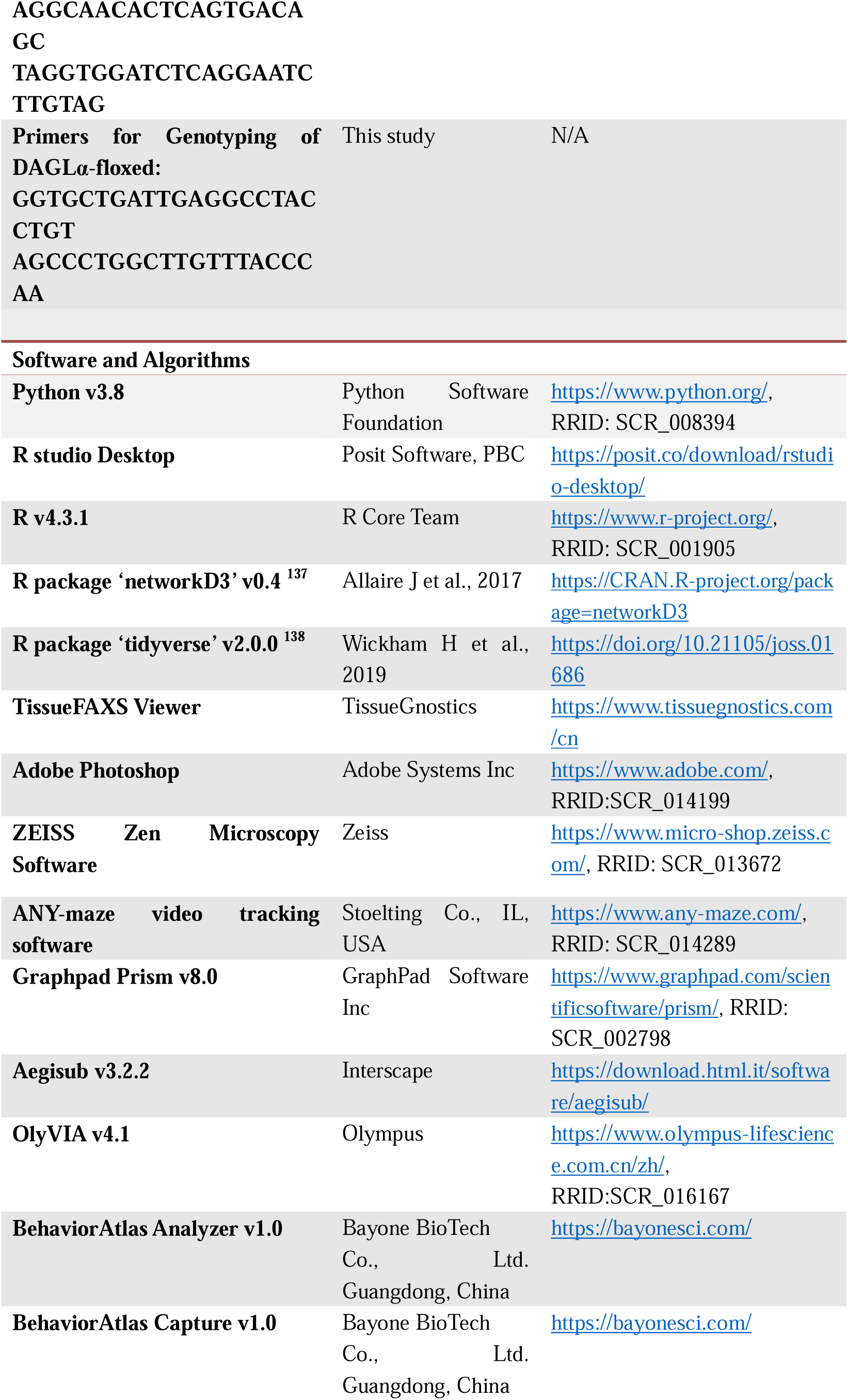
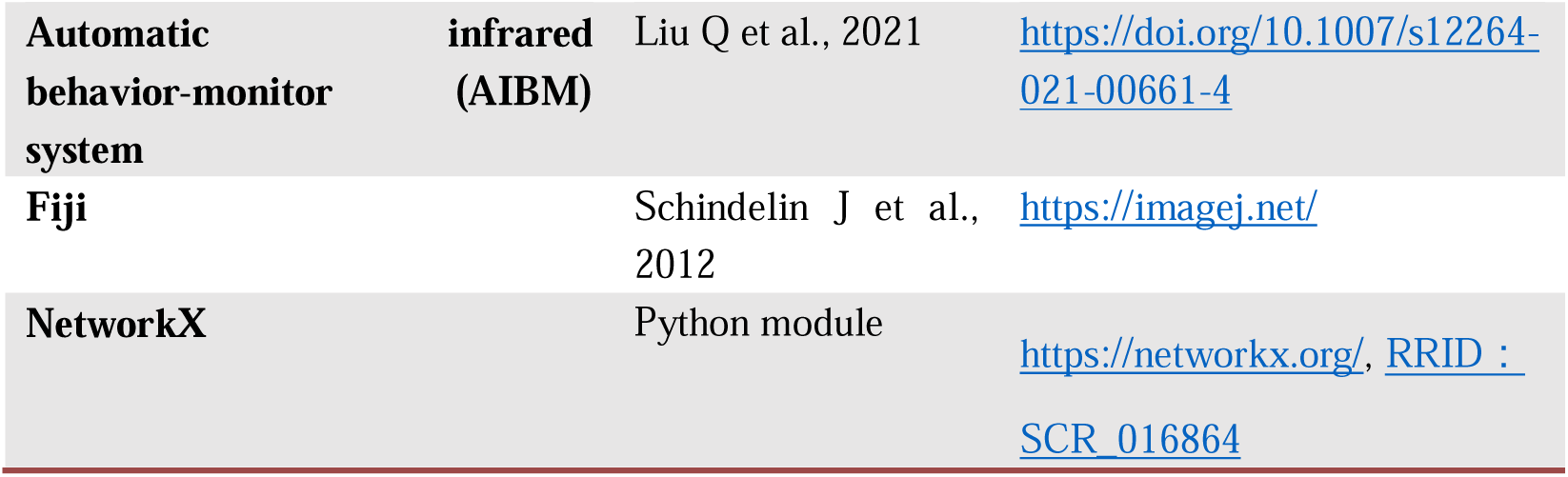

## RESOURCE AVAILABILITY

### Lead contact

Further information and requests for resources and reagents should be directed to and will be fulfilled by the lead contact, Feng Wang.

### Materials availability

This study did not generate new unique reagents.

### Data and code availability

- The data supporting the current study’s findings are available at Zenodo at https://zenodo.org/uploads/16351971 and are publicly available as of the date of publication. Original videos will be shared by the lead contact upon request.
- All original code has been deposited at GitHub at https://github.com/Feng-Wang-Research-Group/A-neural-circuit-for-female-specific-defensive-homeostasis-in-risk-assessment and is publicly available as of the date of publication.
- Any additional information required to reanalyze the data reported in this paper is available from the lead contact upon request.

## EXPERIMENTAL MODEL AND SUBJECT DETAILS

### Animals

All experimental procedures adhered to the guidelines established by the Chinese Council on Animal Care and were approved by the Institutional Animal Care and Use Committee (IACUC) of the Shenzhen Institutes of Advanced Technology (SIAT), Chinese Academy of Sciences (No. SIAT-IACUC-200319-NS-WF-A1162 and No. SIAT-BSI-IRB-240613-NS-WF-A0048). The study utilized animals virgin male and female C57BL/6J and mutant mice aged 8∼16 weeks and virgin male Sprague-Dawley (SD) rats at an age of 12∼20 weeks. All C57BL/6J mice and SD rats were obtained from Zhejiang Vital River Laboratory Animal Technology Co., Ltd (Zhejiang, China). CB_1_R-floxed mice were generous gifts from department of Anesthesiology and Perioperative Medicine, Xijing Hospital, Air Force Medical University, Xi’an, China. The CB_1_R-iCre-EGFP mouse line was generated as described previously^139^. DAGLα-floxed mice were generated specifically for this study by Biocytogen Pharmaceuticals Co., Ltd., Beijing, China in this study. Genotyping was performed using primers listed in key resources table with correct insertion verified through *in situ* hybridization targeting for gene RNA. Mice were housed in a controlled environment at 22–25°C on a 12 h light/12 h dark cycle with *ad libitum* access to food and water. Behavioral tests and recordings were conducted during the 8:00∼16:00.

### Viruses

1. ***In vivo* photometry recording experiments**, 300 nL/site AAV-EF1α-DIO-GCaMP6s (2×10^12^ genomic copies/mL; packaged by Taitool Bioscience Co., Ltd., Shanghai) or AAV-EF1α-DIO-EGFP (3×10^12^ genomic copies/mL; packaged by Fuqiang Xu’s lab at the SIAT) was injected into the unilateral SC (AP −3.8 mm, ML −0.8 mm, DV −1.8 mm from bregma) of CB_1_R-iCre-EGFP mice at a rate of 100 nL/min. 200 nL/site AAV9-GfaABC1D-eCB3.0 (5.46×10^12^ genomic copies/mL; Yulong Li’s Lab at the Peking University) and AAV9-hSyn-RCaMP3 (6×10^14^ genomic copies/mL; packaged by Vigene Biosciences Co., Ltd., Shangdong) was infused into the unilateral LHb (AP −1.85 mm, ML −0.55 mm, DV −2.55 mm from bregma) of adult C57BL/6J mice at a rate of 70 nL/min. 50 nL/site AAV-EF1α-iGABASnFR was infused into the unilateral LHb (AP −1.85 mm, ML, −0.55 mm, DV −2.55 mm from bregma) of CB_1_R-floxed mice at a rate of 70 nL/min.
2. ***CB_1_R* conditional knockout in Dlx5/6^+^ neurons and control experiments**, 300 nL/site AAV-Dlx5/6-GFP or AAV-Dlx5/6-Cre-GFP (5×10^12^ genomic copies/mL; packaged by Liping Wang’s Lab at SIAT) was injected into the bilateral SC (AP −3.8 mm, ML ±0.8 mm, DV −1.8 mm from bregma) at a rate of 100 nL/min. Cre recombination was successful in the target neurons, resulting Cre recombinase expression and the absence of CB_1_R mRNA. (3) **Optogenetic manipulation of SC CB_1_R^+^ Dlx5/6^+^ projections to LHb, ZI, or LP**, 300 nL/site Cre dependent recombinant AAV constructs containing hChR2(H134R)-mCherry (AAV2/9-MDLX-DIO-hChR2(H134R)-mCherry-WPRE-hGH pA; 5×10^12^ genomic copies/ml; packaged by BrainVTA Co., Ltd., Wuhan) or control viruses (AAV2/9-Dlx5/6-DIO-mCherry-WPRE-hGH pA; 5×10^12^ genomic copies/ml; packaged by BrainVTA Co., Ltd., Wuhan) were injected into the unilateral SC (AP −3.8 mm, ML −0.8 mm, DV −1.8 mm from bregma) of CB_1_R-iCre-EGFP mice at a rate of 100 nL/min. (4) **Confirming SC CB_1_R^+^ terminals in the LHb**, 300 nL/site AAV-Dlx5/6-DIO-mCherry and AAV-mDlx5/6-FLEX-tdTomato-T2A-Synaptophysin-EGFP (packaged by Brain Case Co., Ltd., Shenzhen, title: 5×10^12^ genomic copies/mL) were injected into the unilateral SC of CB_1_R-iCre-EGFP mice at a rate of 100 nL/min, respectively. (5) **Discriminating CB_1_R^+^ and CB_1_R^-^ projections in SC Dlx5/6^+^ neurons**, a 1:1 mixture of viruses AAV2/9-Dlx5/6-Cre on-ChR2-mCherry and AAV2/9-Dlx5/6-Cre out-ChR2-YFP (packaged by Shanghai Sunbio Medical Biotechnology Co., Ltd., Shanghai, title: 5×10^12^ genomic copies/mL) was injected into the SC of CB_1_R-iCre-EGFP mice at a rate of 100 nL/min. (6) **Chemogenetic manipulation of LHb CB_1_R^+^ GABAergic projections from the SC**, 300 nL/site Cre dependent recombinant AAV constructs containing hM4D(Gi)-mCherry (rAAV-mDlx5/6-DIO-hM4D(Gi)-mCherry-WPRE-hGH pA; 5×10^12^ genomic copies/ml; packaged by BrainVTA Co., Ltd., Wuhan) or control viruses (AAV2/9-Dlx5/6-DIO-mCherry-WPRE-hGH pA; 5×10^12^ genomic copies/ml; packaged by BrainVTA Co., Ltd., WuhanLtd., Wuhan) were injected into the bilateral SC of CB_1_R-iCre-EGFP mice at a rate of 100 nL/min. (7) **DAGL**α **knock-out and control experiments**, 50 nL/site AAV-EF1α-GFP or AAV-EF1α-Cre-GFP (5×10^12^ genomic copies/mL; packaged by Liping Wang’s Lab at SIAT) was injected into the bilateral LHb (AP −1.85 mm, ML ±0.50 mm, DV −2.75 mm from bregma) at a rate of 70 nL/min.

## METHOD DETAILS

### Experimental design and randomization

For all between-subject analysis, mice were randomly assigned to experimental conditions using cage-level randomization (2-3 littermates split between control and experimental groups) to account for litter effects while minimizing cage-related variance. Experimental replication was achieved through cohort stratification, with conditional CB_1_R knock out models requiring ≥3 batches (n=3-6/ batches) and other behavioral and assays experiments requiring ≥2 batches. For 3D behavior analyses, conditions were randomized, and experimenters remained blinded to group assignments until statistical was completed. Spatial *FISH* experiments employed triple-masking protocols during both experimental execution and computational processing. In neurocircuitry mapping, fiber and cannula placements experiments, histological verification was performed to assess viral transduction efficiency and implant localization. Subjects with low transduction efficiency <50% or deviation from target coordinated were systematically excluded prior to data processing.

### Stereotaxic surgeries

Stereotaxic injection was performed following the protocol outlined in our published work^140^. Mice were anesthetized with pentobarbital (i.p., 80 mg/kg) and positioned on a stereotaxic frame (RWD, Shenzhen, China). Erythromycin eye ointment was applied to protect the eyes. The scalp was shaved, and a ∼1 cm incision was made to expose the cranium. Connective and muscle tissue were carefully removed, and the skull surface was leveled to ensure the height difference between bregma and lambda was <0.05 mm and aligned in the medial-lateral plane. The skull above the target brain region was thinned using a dental drill and carefully removed. Viral injections were performed using a 10-μL Hamilton syringe connected to a 33-gauge needle and a microsyringe pump (UMP3/Micro4, World Precision Instruments, Sarasota, USA). The syringe was filled with mineral oil prior to use. At the injection, the needle remained in place for approximately 5 min to prevent fluid from reflux. Following surgery and a recovery period of at least 7 days, behavioral tests were conducted.

### Cannula implantation

For chemogenetics manipulation and pharmacological experiments, a stainless-steel bilateral cannula (O.D.0.41 mm×I.D.0.25 mm, c.c 1.0 mm, C 2.8 mm, G1 0.5 mm, G2 0.5 mm, RWD Life Science, Shenzhen, China) was implanted into the LHb. For pharmacological experiment, a bilateral cannula (O.D.0.48 mm×I.D.0.34 mm, c.c 2.4 mm, C 4.5 mm, G1 0.5 mm, G2 0.5 mm, RWD Life Science, Shenzhen, China) was implanted in the LHb (coordinates of AP: −1.85Lmm, ML:L±L0.5Lmm, DV: −2.55Lmm), ZI (coordinates of AP: −2.03Lmm, ML:L±L1.2Lmm, DV: −4.4Lmm) or a singular cannula (0.48 mm in diameter, 5 mm in length, RWD Life Science, Shenzhen, China) was implanted in the Re (coordinates of AP: −0.59Lmm, ML:L0.1Lmm, DV: −4.2Lmm). Behavioral tests were performed one week later after implantation.

### Drug delivery

For chemogenetic inhibition of SC-LHb CB_1_R^+^ Dlx5/6^+^ projections in freely behaving mice, Clozapine N-oxide (CNO, Tocris, Cat #:4936) was prepared by dissolved it in vehicle solution (sterile saline with 0.2% DMSO). Aliquots were frozen and freshly diluted with saline to a final concentration of 3 μg/μL before bilaterally administrated into the LHb via a bilateral guide cannula. The cannula was connected to two Hamilton syringes via a polyethylene catheter. For each side of the LHb, 50 nL of CNO or vehicle solution was infused slowly at a rate of 70 nL/min using two syringe pumps. After infusion, mice were habituated for additional 5 min to allow CNO diffusion before the needle was withdrawn. Animals were observed for 30 min prior to behavioral tests.

For pharmacological experiments, the neutral CB_1_R antagonist NESS0327 (Cayman Chemical, 10004184, Michigan, USA) was locally administered to the LHb via a guide cannula. NESS0327 was suspended in saline containing 0.2% Tween 80. On the day of the experiment, a stainless-steel injection cannula (o.d.:0.41; i.d.:0.25, RWD, Shenzhen, China) was inserted into the LHb. The injection cannula, connected to a micro-injection system housing a 10 μL Hamilton syringe via flexible polyethylene tubing, extended 0.4 mm beyond the guide cannula tip. NESS0327 (1 μM/L) was bilateral injected in the LHb at a volume of 50 nL/site. Animals were observed for 3 min before undergoing behavioral tests.

### Histology

#### Fluorescence In Situ Hybridization (FISH)

FISH was carried out following a protocol similar to that previously described^140^. Complementary DNAs (cDNAs) of *CB_1_R*, *GAD2*, *VGLUT2*, or *c-Fos* (sequences from Allen Brain Atlas: CB_1_R, NCBI Accession: NM_007726.2; GAD2, NCBI Accession: NM_008078.1; VGLUT2, NCBI Accession: NM_080853.2; c-Fos, NCBI Accession: NM_010234.2) were cloned from a mouse cDNA library into a pBluescript SK cloning vector. Sequencing was performed to verify the accuracy of the cloned sequences. Antisense complementary RNA (cRNA) probes were synthesized by *in vitro* transcription and labeled with digoxigenin. The following primer sequences were used for cloning of CB_1_R, GAD2, VGLUT2, and c-Fos:

CB_1_R (forward/reverse): GTTCAAGGAGAACGAGGACAAC/CAGCAGGCAGAGCATACTACAG

GAD2 (forward/reverse): TCTTTTCTCCTGGTGGCG/TTGAGAGGCGGCTCATTC

VGLUT2 (forward/reverse): CCAAATCTTACGGTGCTACCTC/TAGCCATCTTTCCTGTTCCACT

c-Fos (forward/reverse): GCCCACCGACCTGCCTGCAA/CTTTCGGTGGGCAGCGGCAC

### Spatial FISH

The specific probes for target RNA (Table S2) were designed by Spatial FISH Ltd (Shenzhen, China). SC sections from adult male (n=3) and female (n=3) C57BL/6J mice were fixed by 4% paraformaldehyde (PFA) and placed within a reaction chamber for subsequent processing. After dehydration and denaturation with methanol, hybridization buffer containing the specific probes were added to the chamber and incubated overnight at 37°C. The samples were then washed three times with PBST, followed by ligation of the target probes in a ligation mix at 25°C for 3 h. After another three washes with PBST, rolling circle amplification was performed using Phi29 DNA polymerase at 30°C overnight. Fluorescent detection probes in hybridization buffer were then applied to the samples. Finally, the samples were dehydrated using an ethanol series and mounted with mounting medium. Images were captured using Leica THUNDER Imaging Systems, and signal dots were decoded to analyze RNA spatial position.

### Immunohistochemistry

Mice were perfused with 20 ml of 1×PBS followed by 20 ml of 4% PFA. Brains were harvested, post-fixed in 4% PFA overnight at 4°C, and dehydrated in 30% sucrose for 48 hours. Using a cryostat (CM1950, Leica, Wetzlar, Germany), 40 μm sections were prepared. Free-floating brain sections were rinsed with PBS (3×5 min), PBST (0.3% Triton X-100 in PBS, 1×20 minutes), and then blocked with 3% normal donkey serum (Solarbio, Cat #:SL050) at room temperature. Sections were incubated overnight at 4°C with primary antibody, including rabbit anti-RFP (1:500 dilution, Rockland, Cat #:600-401-379), rabbit anti-Cre (1:500 dilution, Abcam, Cat #: ab190177), or goat anti-GFP (1:500 dilution, Rockland, Cat #: 600-101-215). After washing in PBST (3×5 min), sections were incubated at room temperature for 1 h with secondary antibodies: donkey anti-rabbit 594 antibody (1:500 dilution, Jackson ImmunoResearch, Cat #:711-585-152), donkey anti-rabbit 488 antibody (1:500 dilution, Jackson ImmunoResearch, Cat #:711-547-003) or donkey anti-goat 488 antibody (1:500 dilution, Jackson ImmunoResearch, Cat #:705-545-003). This was followed by three 10-min washes in 0.01 mol/L PBS. Finally, sections were mounted and covered with the anti-fade reagent DAPI (1:5000 dilution, Sigma-Aldrich, Cat #: D9542).

### Microscopy Imaging and Quantification of Input Neurons

Virus-labeled signals and immunofluorescence-positive cells were traced using an Olympus VS120 or VS200 microscope equipped with a 10× objective and analyzed with Adobe Photoshop software (Adobe, San Jose, USA). For classical image presentation, sections were photographed using a Carl Zeiss ApoTome.2 microscope (Zeiss, Oberkochen, Germany) and processed with Zen software (Zeiss) or TissueFAXS Imaging software (TissueGenostics, Austria) using a 20× objective with tiled imaging (stitched from multiple images). Cell counting was performed manually on every third 40 μm thick section, ensuring a 120 μm intervals between successive sections in the coronal plane. Brain regions were identified based on *The Mouse Brain* (Stereotaxic Coordinates, fourth edition, Franklin, 2013).

### Axon projection mapping

To analyze the projection patterns of SC CB_1_R^+^ Dlx5/6^+^ neurons, every third brain sections (40μm thick) spanning the whole brain was collected. Sections were fully washed with PBS, mounted, and cover-slipped using 50% glycerol. Images were captured using an Olympus microscopy with a 10× objective (Olympus VS200). Brain regions were identified using *The Mouse Brain* (Stereotaxic Coordinates, fourth edition, Franklin, 2013), and projection patterns were analyzed using ImageJ (ImageJ, RRID: SCR_003070). In detail, for each animal, all images were thresholded to remove background noise. Then, 2-5 regions of interest (ROIs) were selected for each brain region, and the average pixel value of each ROI were calculated after background subtraction (fluorescence density excluding fiber-free regions). For each animal, the average fluorescence density of a brain region was determined by averaging the values across all ROIs. Relative density per region was expressed as a fraction of the total density (summed across all regions). Data processing was conducted using the tidyverse R package (v2.0.0), while interactive forced-directed graphs of relative density averages across neuroanatomical regions were generated using the networkD3 package (v0.4).

### Behavioral assays

#### Looming behavior

To assess the defensive behavior of male and female mice in response to life-threatening stimuli, we performed 2D looming behavioral tests using an innately aversive overhead expanding spot (looming task, as previously described^141^). On Day 1, animals were acclimated to the test arena for 10 min. The arena consisted of an open-top acrylic cylinder (“open field”, 50-cm diameter) connected to an adjacent alley (“refuge”, 50 cm×10 cm), allowing free access between compartments. The entire setup was enclosed by a 30-cm high wall (Figure 1B). On Day 2, mice were allowed 5 minutes of free exploration in the open-field arena before being subjected to 1-2 looming trials, depending on the timing accuracy of the first-stimulus, with a 3-minute inter-stimulus interval. The looming stimuli was triggered automatically when mice entered a concentric circular zone (central area: 25 cm diameter) within the arena periphery. The looming stimuli consist of a dark disk expanding from 2° to 40° within 300 ms, maintaining this size for 50 ms before disappearing, and repeating 15 times at 30-ms intervals.

For the SC CB_1_R knock-out experiments, a 3D looming behavioral tests was assessed using multi-view motion-capture device system, as previously described^30^. A transparent looming arena with the same parameter as the 2D behavioral tests was placed at the center of a supporting framework, equipped with four depth cameras (Inter, RealSense D435i) connected to the BehaviorAltas Capture system (version 1.01, Shenzhen Bayone Biotech Co.), ensuring full-view motion capture. Two monitors, mounted on a fixed shelf 45 cm above the bottom of the looming box, delivered uniform and stable white light for behavioral tests. Light intensity was set to 120 lux at the center of the circular trigger zone and 30 lux at the refuge midpoint, matching the brightness parameters of 2D looming behavior setup. Behavioral recordings lasted 15 min, capturing 30 frames per second. matching the brightness parameters of 2D looming behavior setup. The other experimental parameters were consistent with those used in the 2D tests.

### Open field test (OFT) in the multi-view motion-capture device

The open field apparatus (50×50×50 cm, matte white polycarbonate) was calibrated to access anxiety-like behavior and locomotor activity. To collect unobstructed behavioral data for elaborate analysis, a multi-view video capture device was employed to record spontaneous behavior (Figure 6A)^51^. Briefly, a 50×50 cm transparent acrylic square enclosure with white ground was placed at the center of a support framework either 130×130×90 cm (Figure 4L) or a 90×90×75 cm (Figure 6).

Four cameras were rigidly mounted on structural support pillars, with pan-tilt heads finely adjusted to achieve full coverage of the arena. Behavioral recordings were synchronized (30 Hz, 640×480 or 960×540-pixel resolution) and processed using BehaviorAtlas Capture Software® v2.0.0.

### 3D Behavioral data processing

Data acquired from the BehaviorAtlas Capture System underwent three sequential stages as previously described^51^: unsupervised movement annotation, supervised label refinement with manual correction of misclassified movement labels, and ethological quantification.

### Unsupervised movement annotation pipeline

Using pre-training body point data from images, the skeleton of each mouse included in the study was automatically reconstructed by BehaviorAltas Analyzer software, incorporating camera calibration information. Movements were classified by clustering skeleton kinematic features for each frame and assigning digital labels.

### Supervised movement labeling

Movements with postural similarities, particularly those occurring in low-light refuge areas, were subjected to dual-criteria validation with skeleton kinematic parameters such as speed, head-to-limb angle, body length, and distances between the nose and limbs. Specific movements like grooming, pausing, and looking up were systematically reviewed and reclassified when discrepancies arose between 3D kinematic data and manual annotations (see below). Additionally, movements such as flight and freezing were manually annotated by experts due to significant inter-individual variability in kinematic patterns.

Kinematic reclassification protocol

| **Movement** | **Threshold** |
| --- | --- |
| Pausing | Average velocity (based on back coordinate) < 30 mm/s and the movement duration > 2 s |
| Hunching | Average nose height ranges from 30 to 45 mm and average back height > 25 mm |
| Looking up | Average nose height > 20mm and average front claw height < 15 mm |
| Right turning | Average velocity (based on back coordinate) > 50 mm/s and body angle from top-view (neck-back-root tail) >15° |
| Grooming | Average body angle in 3D space (neck-back-root tail) < 120° |
| Rotating | Average velocity (based on back coordinate) > 30 mm/s and body angle from top-view (neck-back-root tail) >20° |

### Ethograms

Four key defensive behavior metrics, flight probability, response latency, return time, and time in the refuge, were quantified through manual frame-by-frame analysis (Figure 2D-E). Hierarchical ethograms were constructed by segmenting movement fragments within specific time windows (Figure 2H-I). Trajectory analysis was performed to quantify mouse locomotion by tracking the coordinates of the mouse’s back, integrating spatial and temporal movement parameters. Defensive behavior quantification curves were created by mapping three stereotypic responses, risk-assessment posturing, flight initiation, freezing duration, against threat-exposure timeline, with group-averaged movement probability on the y-axis and time on the x-axis. Peak response markers indicated behavior-specific latency (Figure 2L-M). Defensive behavior analysis also incorporated nasal elevation dynamics and locomotor velocity, derived from 3D pose estimation outputs (Figure 2N-O).

### Sample classification and segregation

Movement classification was performed using a random forest classifier with PCA-reduced kinematic features, encompassing all movements and their frequencies. Random forest machine learning algorithms were applied to movement data for group classification, with accuracy evaluated through detailed confusion matrices.

### Movement transition and central node definition

Movement transitions were analyzed using probability matrices that captured how often one movement followed another, disregarding the duration of each movement. Line thickness in the diagrams represented the frequency of transitions, while dot size indicated the likelihood of specific movement events (Figure S6I-L). Transitions with probabilities below 0.05, after statistical adjustments, were excluded. Network diagrams were generated using Python’s NetworkX package (v2.8.4) with spring layout settings.

### Elevated plus maze (EPM)

The EPM test was performed a using maze made of white acrylic glass, comprising of a central platform (5×5 cm), two opposing open arms (32.5×5 cm), and two opposing closed arms (32.5×5×32.5 cm). The maze was elevated 50 cm above the floor. Mice were placed at the junction of the open and closed arms, facing the closed arms. Behavioral tracking (5-minute sessions) was conducted using ANY-maze® v7.4 software, with body centroid tracking. Time spent in each area (center, open arms, closed arms) was quantified.

Between behavior trials, the arena was cleaned by scrubbing with a 25% ethanol solution.

### Chronic rat stress model

The experimental protocol was modified from previous studies^142,143^. In the chronic stress model, mice were exposed to a rat for 10 consecutive days. Each day, a mouse was placed in a cage containing a rat for 5 seconds and then co-housed with the rat using a ventilated plexiglass divider. This apparatus allowed odor transmission while preventing physical contact, ensuring continuous exposure to predator odors. Behavioral tests, including the open-field test and elevated-plus maze, were conducted at two time points: 3 days prior to the establishment of the model (Pre-stress) and on day 10 post-exposure to rat-induced stress (Post-stress).

### Patch-clamp electrophysiology

Coronal brain slices (300 μm thick) containing the SC region (bregma 3.2–4.4 mm) were prepared from CB_1_R-iCre-EGFP mice using standard procedures. Brains were quickly separated and chilled in ice-cold modified artificial cerebrospinal fluid (ACSF) containing (in mM): 110 Choline Chloride, 2.5 KCl, 1.3 NaH_2_PO_4_, 25 NaHCO_3_, 1.3 Na-Ascorbate, 0.6 Na-Pyruvate, 10 Glucose, 2 CaCl_2_, 1.3 MgCl_2_. SC slices were sectioned in ice-cold modified ACSF using a Leica vibroslicer (VT-1200S). Following sectioning, slices were allowed to recover for 30 min in a storage chamber containing regular ACSF at 32∼34 °C (in mM): 125 NaCl, 2.5 KCl, 1.3 NaH_2_PO_4_, 25 NaHCO3, 1.3 Na-Ascorbate, 0.6 Na-Pyruvate, 10 Glucose, 2 CaCl_2_, 1.3 MgCl_2_ (pH 7.3∼7.4 when saturated with 95% O2/5% CO2). Slices were then maintained at ∼25 °C until transferred to the recording chamber. All solutions had an osmolarity of 300∼320 mOsm/kg. For electrophysiological experiments, slices were visualized using infrared optics on an upright microscope (Eclipse FN1, Nikon Instruments) with a 40× water-immersion objective. The recording chamber was continuously perfused with oxygenated ACSF (2 ml/min) at 34 °C. Patch pipettes were pulled by a micropipette puller (Sutter P-2000 Micropipette Puller) and had a resistance of 3–5MΩ. Electrodes were filled with intracellular solution containing (in mM): 130 potassium gluconate, 1 EGTA, 10 NaCl, 10 HEPES, 2 MgCl_2_, 0.133 CaCl2, 3.5Mg-ATP, 1 Na-GTP. In SC slices, mCherry-positive neurons were patched in current-clamp mode. Action potentials were elicited by stimulation SC neurons expressing ChR2 with blue light pulse (10 ms duration, 20 Hz).

### Optical fiber implantation

Fifteen min after AAV injection, a ceramic ferrule with an optical fiber (200 μm in diameter, N.A. 0.37) was implanted, positioning the optical fiber tip above the LHb (AP: −1.90 mm, ML: ±0.50 mm, DV: −2.55 mm from bregma), ZI (AP: −1.79 mm, ML: +1.10 mm, DV: −4.3 mm from bregma), or LP (AP: −2.03 mm, ML: +1.20 mm, DV: −2.2 mm from bregma). Optogenetic stimulation was performed 4 weeks after fiber implantation.

Three weeks after AAV injection, a ceramic ferrule with an optical fiber (200 μm diameter, N.A. 0.37) was implanted, positioning the fiber tip above the SC (AP: −3.8 mm, ML: −0.8 mm, DV: −1.5 mm from bregma), or LHb (AP: −1.90 mm, ML: ±0.50 mm, DV: −2.45 mm from bregma). Fiber photometry recording was conducted one week after fiber implantation.

### Optogenetics

Optogenetic manipulations were performed using laser systems with wavelength-specific settings. Terminal activation in SC-LHb CB_1_R^+^Dlx5/6^+^ experiments: a blue laser (473 nm; Aurora-300-470, NEWDOON, Hangzhou) was used to deliver 10 mW/mm² at the fiber tip, with 5-s illumination, 20 Hz frequency, and 10-ms pulse duration.

### Fiber photometry

1. Fluorescence signals of GCaMP6s and GABASnFR were recorded via fiber photometry as previously described^142^, using 470-nm LEDs for target excitation and 410-nm LEDs for isosbestic control. The 410-nm light intensity was calibrated to 20 μW/mm² at the fiber tip to match the 470-nm intensity. (2) Fluorescence signals of eCB3.0 and RCaMP3 were recorded via fiber photometry (vista, Inper), using 470-nm, 560 -nm LEDs for target excitation and 410-nm LEDs for isosbestic control. The 410-nm light intensity (0.5 mW/mm²), 470-nm light intensity (0.8 mW/mm²), 561-nm light intensity (1.2 mW/mm²), was calibrated to at the fiber tip. Behavioral video synchronization was achieved through dedicated system software. Raw data were smoothed using a 5-25-point moving average variance, with variance analyzed across different experiments. ΔF/F, areas under the curve, and peak values were calculated using least-squares regression to scale the 410-nm to 470-nm or 560-nm traces.

## QUANTIFICATION AND STATISTICAL ANALYSIS

### Statistics

For all experiments, statistical analyses were performed using either unpaired two-tailed Student’s t test or a two-way ANOVA followed by Holm-Sidak’s multiple comparisons test. The statistical significance threshold was set at α=0.05, with significance levels indicated as \**p* < 0.05, \*\**p* < 0.01, *** *p* < 0.001, **** *p* < 0.0001). Exact *p*-values are provided in the figure legends and Table S1. The “n” in these analyses represents the number of mice, cells, or brain sections. Statistical analyses were carried out using GraphPad Prism 10 (GraphPad Software Inc.). Additionally, a custom Python script was utilized to evaluate the differences in movement transition and postures through a permutation test (10,000 iterations), comparing mean disparities between the two groups. Comprehensive statistical information is available in the figure legends and Table S1.

---

**Figure S1.**
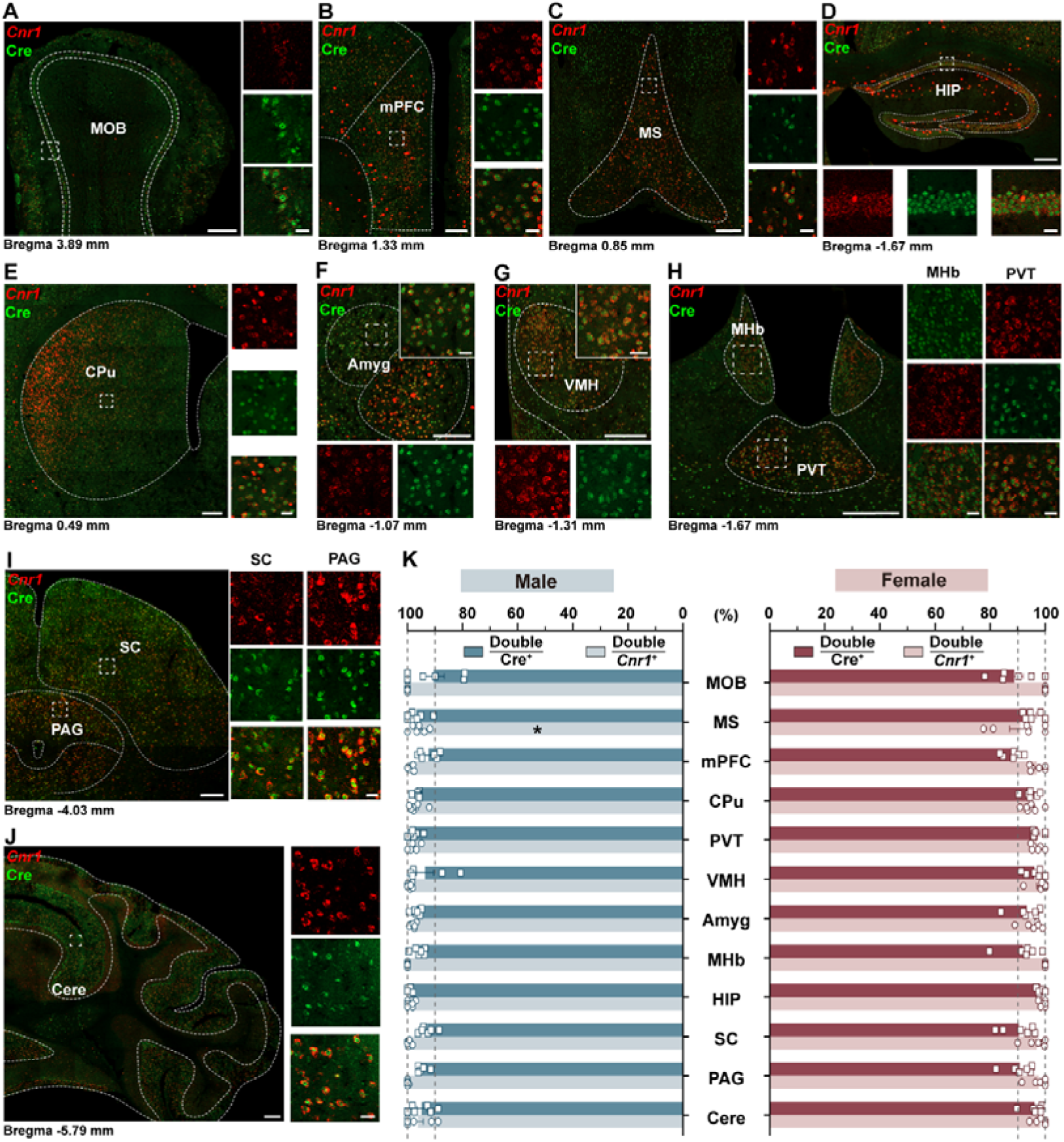
CB_1_R^+^ cells characterization in male and female brains, related to Figure 1. (A)-(I) Diagram representation showing the colocalization of CB_1_R mRNA (red) and Cre protein (green) positive cells in various brain regions of a female mouse, including the MOB (A), mPFC (B), MS (C), HIP (D), CPu (E), Amyg (F), VMH (G), MHb (H), PVT (H), SC (I), PAG (I), Cere (J). Scale bar, 200 μm and 50 μm (enlarged view). (K) Statistical analysis illustrating the percentage of overlap between CB_1_R mRNA (red) and Cre protein (green) positive cells in the mouse brain. Data are presented as mean ± SEM (n = 6 per group). Two-Way ANOVA followed by Sidak’s multiple comparisons post hoc test was performed. \**p*<0.05. For detailed *p* values, refer to Table S1. Abbreviations: MOB, main olfactory bulb; mPFC, medial prefrontal cortex; MS, medial septal nucleus; HIP, hippocampus; CPu, caudate putamen (striatum); Amyg, amygdala; VMH, ventromedial hypothalamus; MHb, medial habenular nucleus; PVT, paraventricular thalamic nucleus; SC, superior colliculus; PAG, periaqueductal gray; Cere, cerebellum.

**Figure S2.**
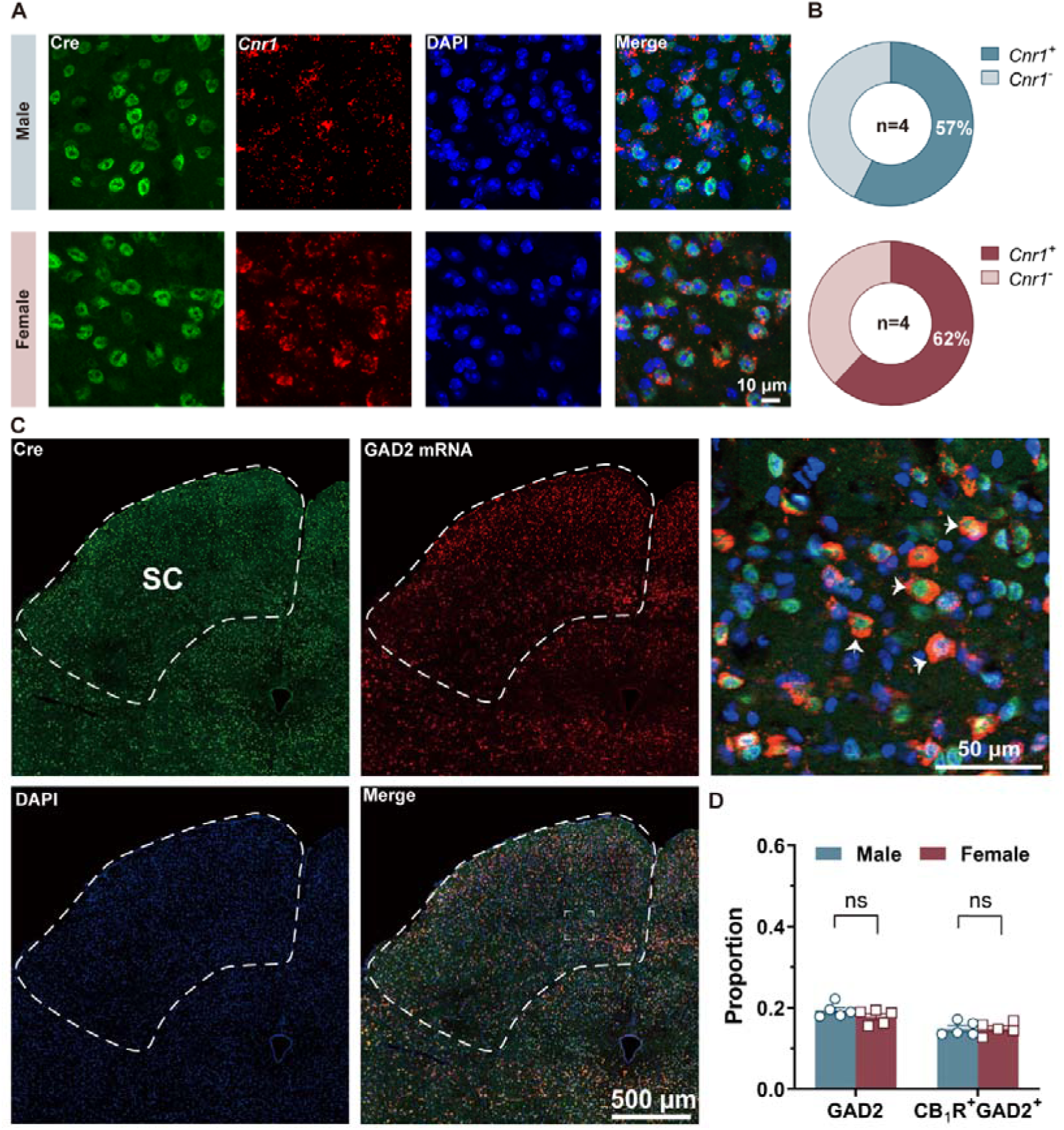
SC neurons majority are CB_1_R^+^ without sex differences, related to Figure 1. (A) Diagram showing the colocalization of CB_1_R mRNA (red) and Cre (green) positive cell in the SC. (B) Pie chart illustrating the proportions of *Cnr1*^+^ and *Cnr1*^-^ cells in the SC for male (top) and female (bottom) mice. Unpaired Student’s t test with the Mann-Whitney test was performed. (C) Representative fluorescence *in situ* hybridization images showing CB_1_R^+^ Cre expression in GAD2 neurons in the SC (Green, Cre; Red, GAD2 mRNA). Scale bar, 500 μm, and 50 μm (enlarged view). White arrows indicate the GAD2 mRNA fluorescent signals co-labeled with CB_1_R positive Cre signals in each panel. (D) Percentage of GAD2^+^ and CB_1_R^+^ GAD2 neurons in males and females. Two-Way ANOVA followed by Sidak’s multiple comparisons post hoc test was performed. Data are presented as means ± SEM, n = 4 per group (B), 5 per group (D). ns (not significant). For detailed *p* values, refer to Table S1.

**Figure S3.**
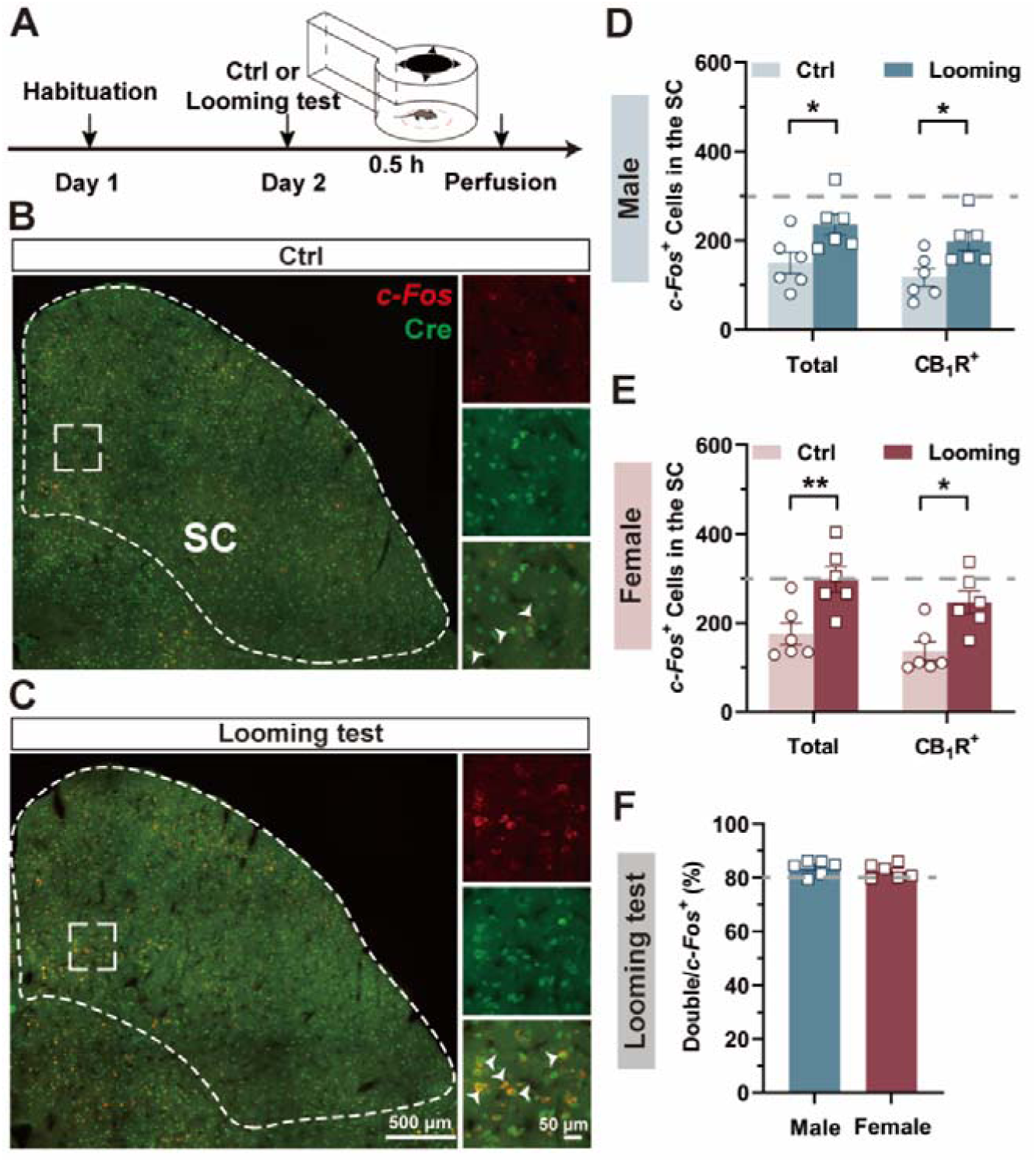
Visual threat activates SC CB_1_R^+^ neurons, related to Figure 1. (A) Timeline of the experimental protocol for looming stimuli and a schematic representation of the paradigm using a refuge-containing open field apparatus. (B-C) Representative images showing c-Fos mRNA expression in the SC under the control condition (B) and after the looming test (C) (red, c-Fos mRNA; green, Cre protein; scale bar, 500 μm and 50 μm (enlarged view). White arrows indicate c-Fos mRNA fluorescent signals co-labeled with CB_1_R-positive Cre signals in each panel. (D-E) Quantification of c-Fos mRNA-labeled neurons and their colocalization with CB_1_R-positive Cre signals in the SC of males (D) and females (E). Two-Way ANOVA followed by Sidak’s multiple comparisons post hoc test was performed. (F) Percentage of CB_1_R^+^ c-Fos mRNA labeled neurons across the total c-Fos mRNA-labeled neurons following the looming test in males and females. Unpaired Student’s t test with Mann-Whitney test was performed. Data are presented as means ± SEM, n=6 per group (D-F). \**p*<0.05; \*\**p*< 0.01. For detailed *p* values, refer to Table S1.

**Figure S4.**
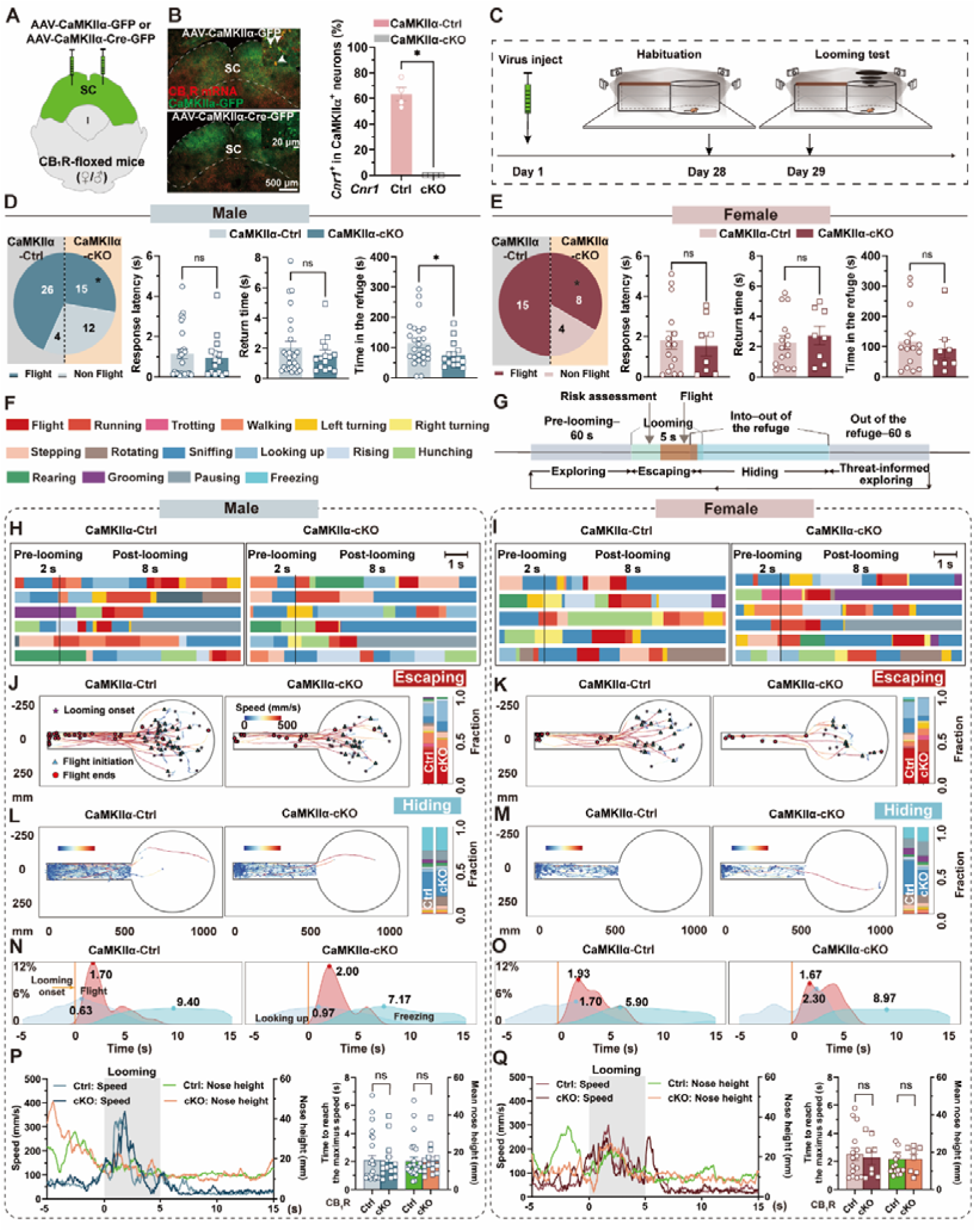
Deletion of CB_1_R in SC CaMKIIα^+^ neurons suppressed defensive behavior in both sexes, related to Figure 2. (A) Schematic representation of viral injection of AAV-CaMKIIα-GFP or AAV-CaMKIIα-Cre-GFP into the bilateral SC of CB_1_R-floxed mice. (B) Left: Representative images showing AAV-CaMKIIα-GFP (top) and AAV-CaMKIIα-Cre-GFP (bottom) expression in the SC, alongside CB_1_R mRNA (red) and CaMKIIα^+^ cells bodies (green) in CaMKIIα-Ctrl and CaMKIIα-cKO group. Scale bar, 500 μm and 20 μm (enlarged view). White arrows indicate CB_1_R mRNA fluorescent signals colocalized with GFP positive signals. Right: Quantification of CB_1_R^+^ cells in CaMKIIα^+^ neurons in CaMKIIα-Ctrl and CaMKIIα-cKO groups for both sexes. (C) Experimental timeline and schematic paradigm of virus injection and the looming test in a refuge-containing apparatus. (D and E) Defensive strategy (left) and flight-to-refuge behaviors (from left to right: response latency, return time, time spent in the refuge) in male (D) and female (E) mice across CaMKIIα-Ctrl and CaMKIIα-cKO group. (F) Sixteen movement types during defensive behavior, represented by corresponding colors. (G) Schematic paradigm of the different stages of defensive behavior: exploring, escaping, hiding and threat-informed exploring. (H and I) Ethograms showing the 16 movement types in males (H) and females (I) across groups (CaMKIIα-Ctrl group on the left, CaMKIIα-cKO group on the right). Scale bar, 1 s. (J-K) Trajectories (left and middle) and quantification of fractions (right) of mice during the escaping stage (from the onset of looming to the end of flight, J, K) and hiding in the refuge (L, M) in males (J, L) and females (K, M) across CaMKIIα-Ctrl and CaMKIIα-cKO groups. Histograms show the percentage of mice occupying horizontal and vertical positions in the apparatus. Symbols indicate mouse locations: five-pointed star for looming onset, triangle for flight initiation, and circle for flight termination. (N and O) Distribution and positions of mice during flight, looking up, and freezing from 5 s pre-looming to 15 s post-looming in males (N) and females (O) across groups (CaMKIIα-Ctrl on the left, CaMKIIα-cKO on the right). (P and Q) Left: Real-time speed and nose height from 5 s pre-looming to 15 s post-looming in males (P) and females (Q) across groups. Right: Quantification of the time taken to reach maximum speed and mean nose height during looming stimuli in males (P) and females (Q). Data are presented as mean ± SEM (B, D, E, P, Q) or mean (J-O). n=4 per group (B); 30 male CaMKIIα-Ctrl, 27 male CaMKIIα-cKO (D), 15 female CaMKIIα-Ctrl, 12 female CaMKIIα-cKO (E); 26 male CaMKIIα-Ctrl and 15 CaMKIIα-cKO, 15 female CaMKIIα-Ctrl and 8 CaMKIIα-cKO (K-R). Two-Way ANOVA with Sidak’s multiple comparisons post hoc test (J-M) and Unpaired Student’s t test with Mann-Whitney test (B, D, E, P, Q) were performed. ns (not significant), * *p*<0.05. For detailed *p* values, refer to Table S1.

**Figure S5.**
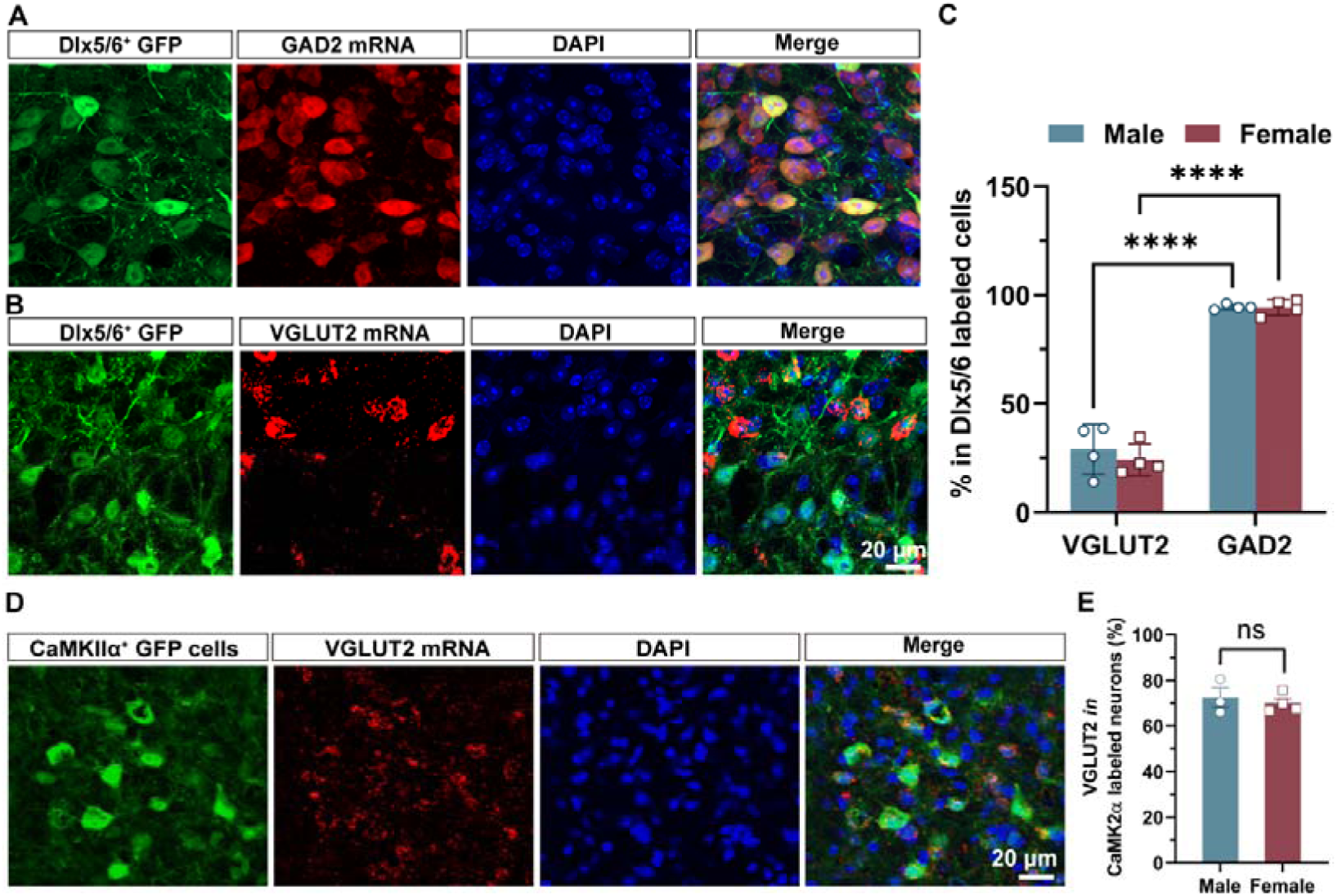
Selective GABAergic targeting with Dlx5/6-driven AAVs, related to Figure 2. (A and B) Representative images showing SC Dlx5/6^+^ GFP signals in GAD2 (A) and VGLUT2 (B) SC neurons. Scale bar, 20 μm. (C) Percentage of SC Dlx5/6^+^ neurons that are positive for GAD2 (A) or VGLUT2 (B) mRNA as determined by *ISH* in males and females. Data are presented as mean ± SEM. (D) Representative image showing CaMKIIα^+^ GFP signals in the VGLUT2 SC neurons (green, CaMKIIα^+^ GFP signals; red, VGLUT2 mRNA; blue, DAPI; scale bars, 20 μm). (E) Percentage of SC CaMKIIα^+^ neurons that are positive for VGLUT2 (E) mRNA as determined by *ISH* in males and females. Data are presented as mean ± SEM. n = 4 per group (C), Two-Way ANOVA with Sidak’s multiple comparisons post hoc test was performed, \*\*\*\**p*<0.0001. n = 3 for males and n=4 for females (E). Unpaired Student’s t test with the Mann-Whitney test. ns (not significant), \**\*\*\*p*<0.0001. For detailed *p* values, refer to Table S1.

**Figure S6.**
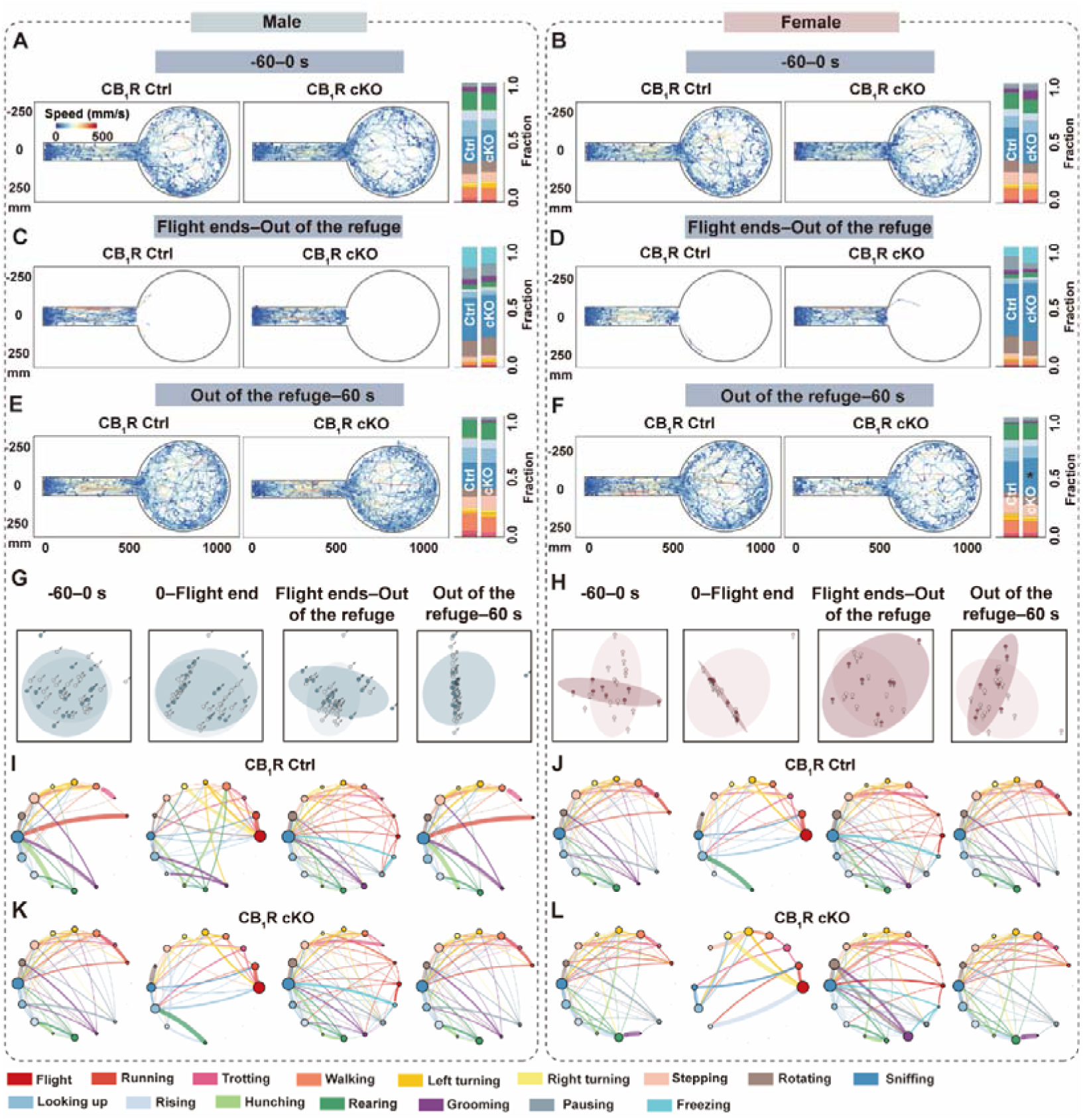
Deletion CB_1_R in SC Dlx5/6^+^ neurons increases female sniffing during threat-informed exploration, related to Figure 2. (A and B) Trajectories (left and middle) and quantification of fractions (right) of mice during the pre-looming stage in males (A) and females (B) across different groups. (C and D) Trajectories (left and middle) and quantification of fractions (right) of mice during the hiding stage in males (C) and females (D) across different groups. (E and F) Trajectories (left and middle) and quantification of fractions (right) of mice during the threat-informed exploring stage in males (E) and females (F) across different groups. Data are presented as mean ± SEM, n=22 (E) male CB_1_R Ctrl, 22 (E) male CB_1_R cKO, 16 (F) female CB_1_R Ctrl, and 16 (F) female CB_1_R cKO. Two-Way ANOVA with Sidak’s multiple comparisons post hoc test was performed. \**p*<0.05. For detailed *p* values, refer to Table S1. (G and H) Behavioral features across different stages (from left to right, pre-looming, escaping, hiding and threat-informed exploring) visualized by embedding 16-dimensional movement fractions into a low-dimensional space using principal component analysis (PCA) for males (G) and females (H). (I-L) Movements transition probabilities across different stages (from left to right, pre-looming, escaping, hiding and threat-informed exploring) for the male CB_1_R Ctrl (I), male CB_1_R cKO (K), female CB_1_R Ctrl (J), and female CB_1_R cKO (L). Each movement is represented as a state, with circle size indicating the number of movement bouts, where larger circles correspond to higher occurrence frequencies.

**Figure S7.**
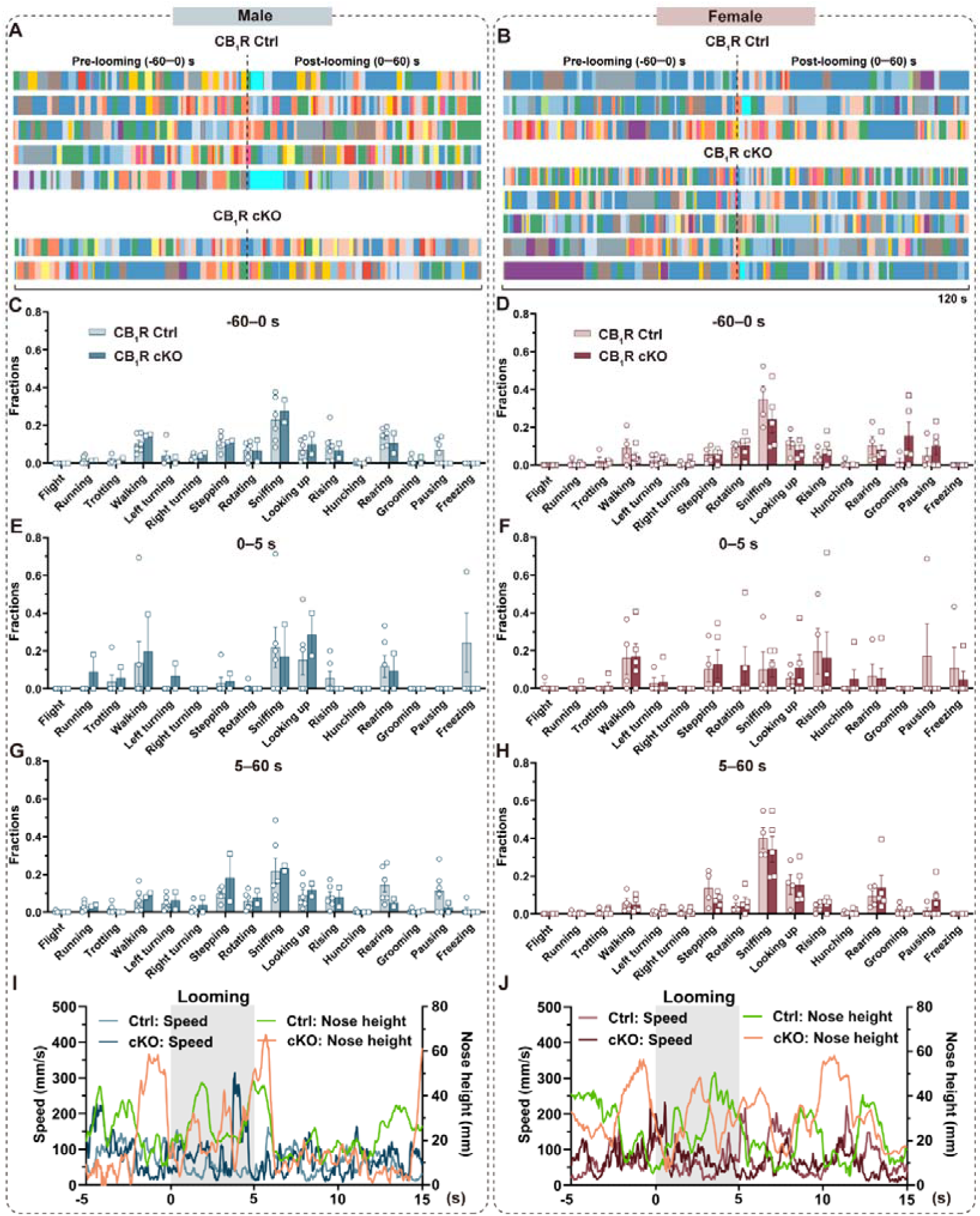
Flight-to-refuge deficits across groups following visual threats, related to Figure 2. (A and B) Ethograms depicting 16 movements from 60 s pre-looming to 60 s post-looming for males (A) and females (B) in the CB_1_R Ctrl (top) and CB_1_R cKO (bottom) group. (C and D) Comparison of movement fractions during the pre-looming stage in males (C) and females (D) across different groups. (E and F) Comparison of movement fractions during the looming stimuli stage in males (E) and females (F) across different groups. (G and H) Comparison of movement fractions during the post-looming stage in males (C) and females (D) across different groups. (I and J) Real-time speed and nose height data from 5 s pre-looming to 15 s post-looming for males (I) and females (J) across different groups. Data are presented as mean ± SEM for C-H and as mean for I, J. n=6 male Ctrl group, 2 male CB_1_R-KO group (C, E, G, I), 4 female CB_1_R Ctrl group, 5 female CB_1_R cKO group (D, F, H, J). For detailed data information, refer to Table S1.

**Figure S8.**
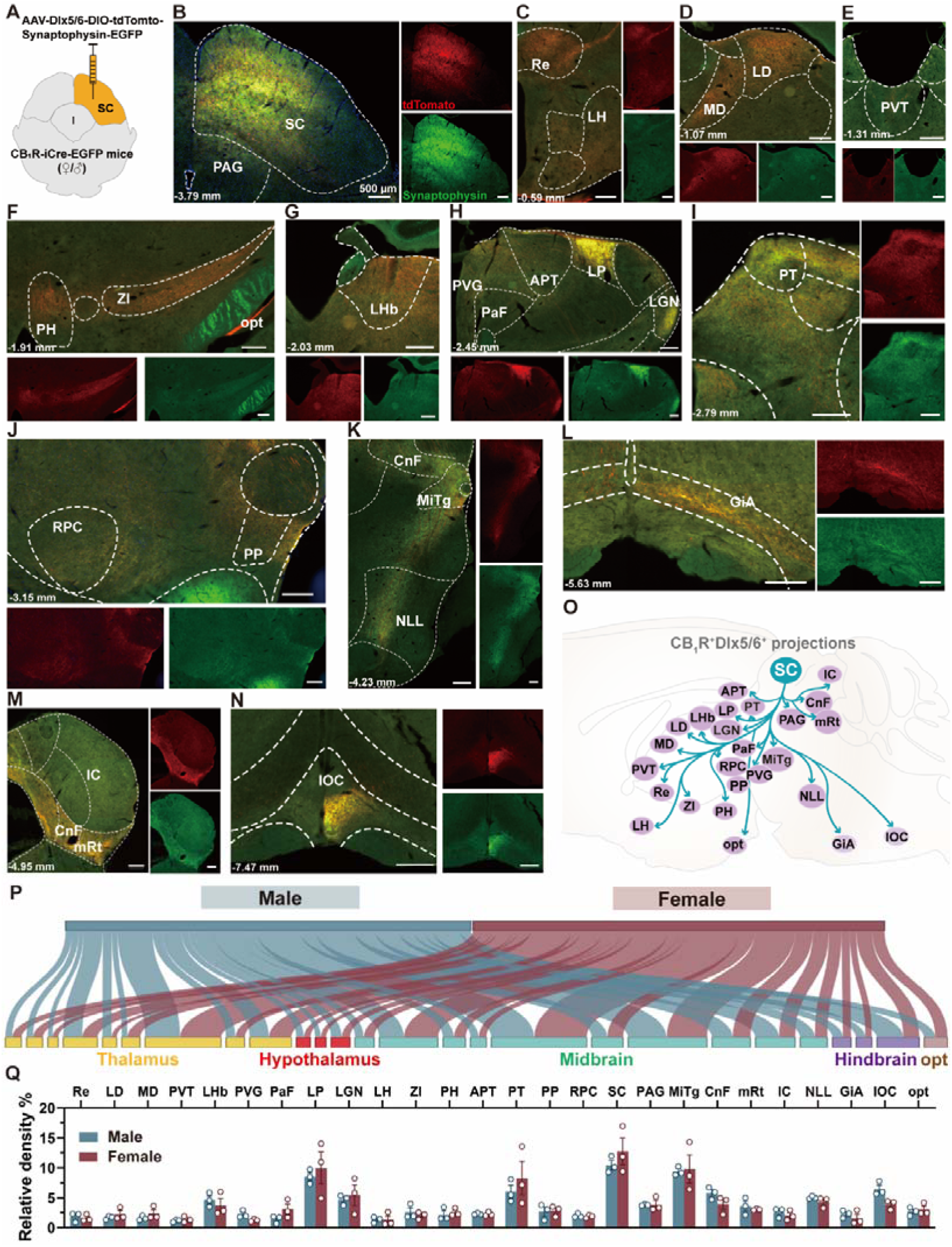
Comparable downstream targeting of SC CB_1_R^+^Dlx5/6^+^ neurons in males and females, related to Figure 3. (A) Schematic illustration of the surgical procedure for anterograde tracing of SC CB_1_R**^+^**Dlx5/6^+^ neurons. (B) Representative image showing viral infection in SC CB_1_R**^+^**Dlx5/6^+^ neurons in a female mouse. Red, CB_1_R**^+^**Dlx5/6^+^ starting cells; green, terminals of CB_1_R**^+^**Dlx5/6^+^ cells. Scale bar, 500 μm. (C-N) Representative image of various brain regions targeted by SC CB_1_R**^+^**Dlx5/6^+^ neurons in a female mouse: Re (C), LH (C), LD (D), MD (D), PVT (E), ZI (F), PH (F), opt (F), LHb (G), PVG (H), PaF (H), APT (H), LP (H), LGN (H), PT (I), RPC (J), PP (J), CnF (K), MiTg (K), NLL (K), IC (L), GiA (L), CnF (M), mRt (M), and IOC (N). Abbreviations: Re, reuniens thalamic nucleus; LH, lateral hypothalamus; LD, laterodorsal thalamus; MD, mediodorsal thalamus, PVT, paraventricular thalamus, LHb, lateral habenula; PVG, periventricular gray; PaF, parafascicular thalamic nucleus; APT, anterior pretectal nucleus, LP, lateral posterior thalamus; LGN, lateral geniculate nucleus; PH, posterior hypothalamus; ZI, zona incerta, opt, optic fiber; PT, pretectal nucleus (PT); RPC, red nucleus, parvicellular part; PP, peripeduncular nucleus; CnF, cuneiform nucleus; MiTg, microcellular tegmental nucleus; NLL, nucleus of the lateral lemniscus pontine reticular nucleus; IC, inferior colliculus, mRt, mesencephalic reticular format; GiA, gigantocellular reticular nucleus, alpha part; and IOC, inferior olive, subnucleus C of medial nucleus. (O) Schematic representation showing outputs of SC CB_1_R**^+^**Dlx5/6^+^ neurons. (P) Sankey network showing projections of SC CB_1_R**^+^**Dlx5/6^+^ neurons in male and female mice, with curve width representing connection strength. (Q) Quantification of normalized fluorescence density in ipsilateral downstream brain subregions targeted by SC CB_1_R**^+^**Dlx5/6 neurons in male and female mice. Data are presented as mean ± SEM. n=3 each group. Two-Way ANOVA with Sidak’s multiple comparisons post hoc test was performed. For detailed *p* values, refer to Table S1.

**Figure S9.**
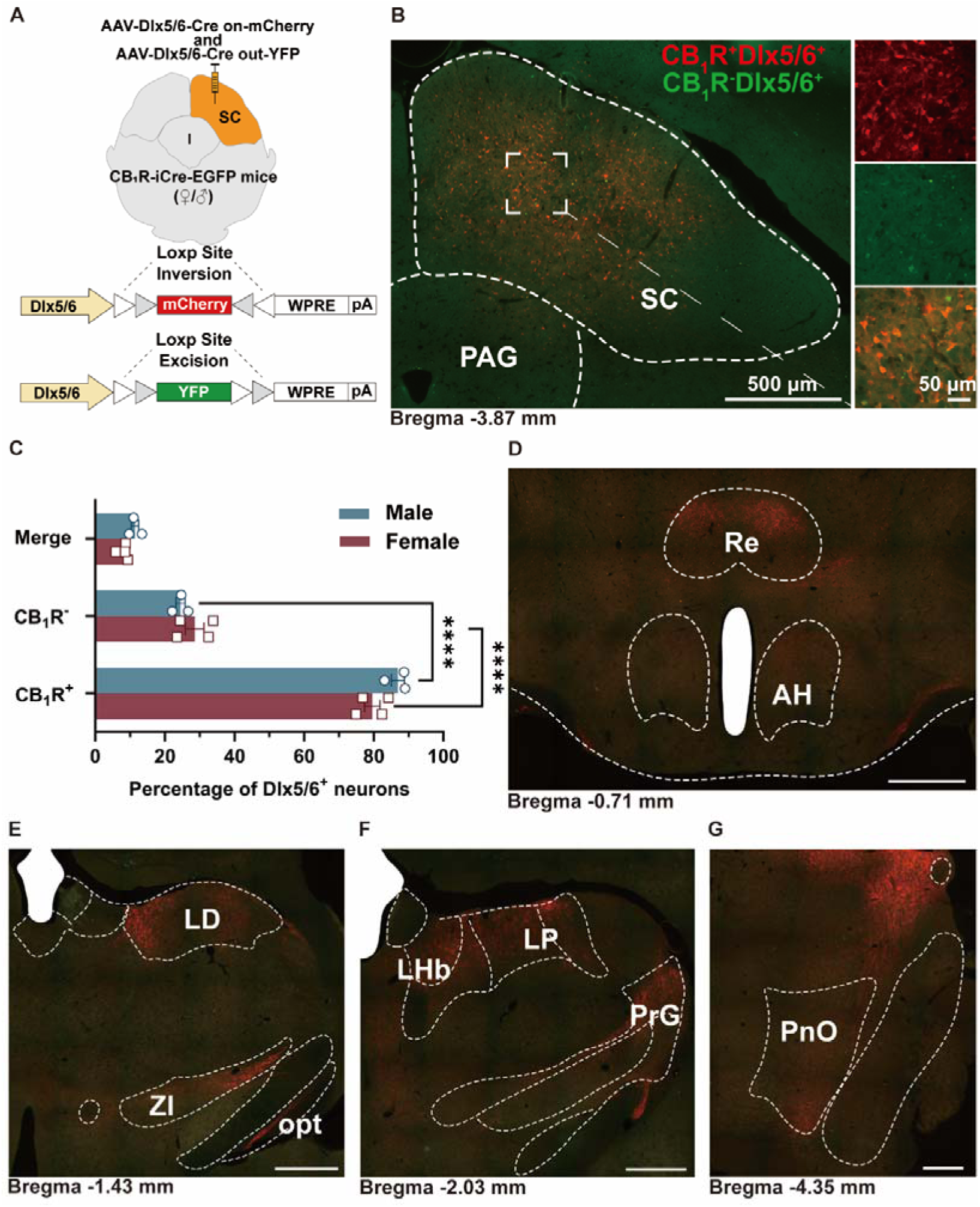
SC CB_1_R^+^Dlx5/6^+^ neurons dominate long-range projections, related to Figure 3. (A) Schematic representation of the virus tracing strategy. (B) Representative fluorescent micrographs of a coronal brain slice showing the injection site in the SC (left) and an enlarged view of the dashed square area (right). Scale bars, 500 μm and 50 μm (enlarged view). (C) Quantification of the percentage of CB_1_R^+^, CB_1_R^-^, and non-specific overlapped signals with mCherry and YFP fluorescence in Dlx5/6^+^ starter neurons. Data are presented as mean ± SEM. n = 3 males, 4 females. Unpaired Student’s t test with the Mann-Whitney test was performed. \*\*\*\**p*< 0.0001. For detailed p values, refer to Table S1. (D-G) Representative images showing downstream projections of SC CB_1_R^+^Dlx5/6^+^ neurons in various brain regions including the reuniens thalamic nucleus (Re), anterior hypothalamus (AH), lateral dorsal thalamus (LD), zona incerta (ZI), optic fiber (opt), lateral habenular (LHb), lateral posterior thalamus (LP), pregeniculate area (PrG) and the pontine reticular nucleus, oral part (PnO).

**Figure S10.**
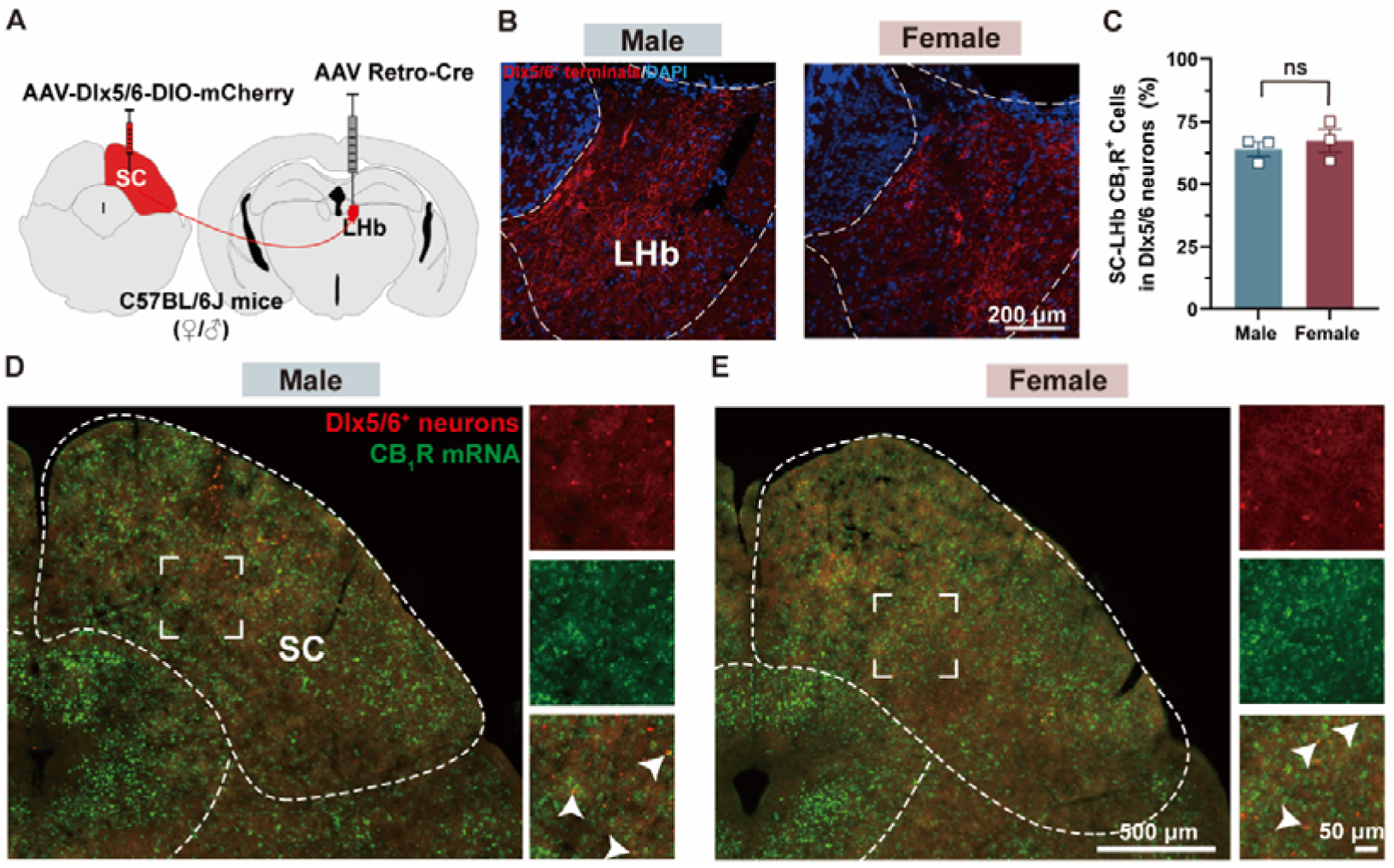
Retrograde tracing identifies CB_1_R^+^Dlx5/6^+^ SC-LHb circuit, related to Figure 3. (A) Schematic representation of the AAV-Retro tracing strategy for SC-LHb projections using unilateral injection into the SC and LHb. (B) Representative fluorescent micrographs showing Dlx5/6^+^ mCherry expression in the LHb. Scale bar, 200 μm. (C) Quantification of the CB_1_R^+^ SC-LHb population within Dlx5/6^+^ neurons. Data are presented as mean ± SEM, n = 3 per group. Unpaired Student’s t test with the Mann-Whitney test was performed. ns (not significant). For detailed *p* values, refer to Table S1. (D and E) FISH analysis showing mCherry expression in the SC co-labeled with CB_1_R mRNA signals in male (D) and female (E) mice. Scale bars, 500 μm and 50 μm (enlarged view).

**Figure S11.**
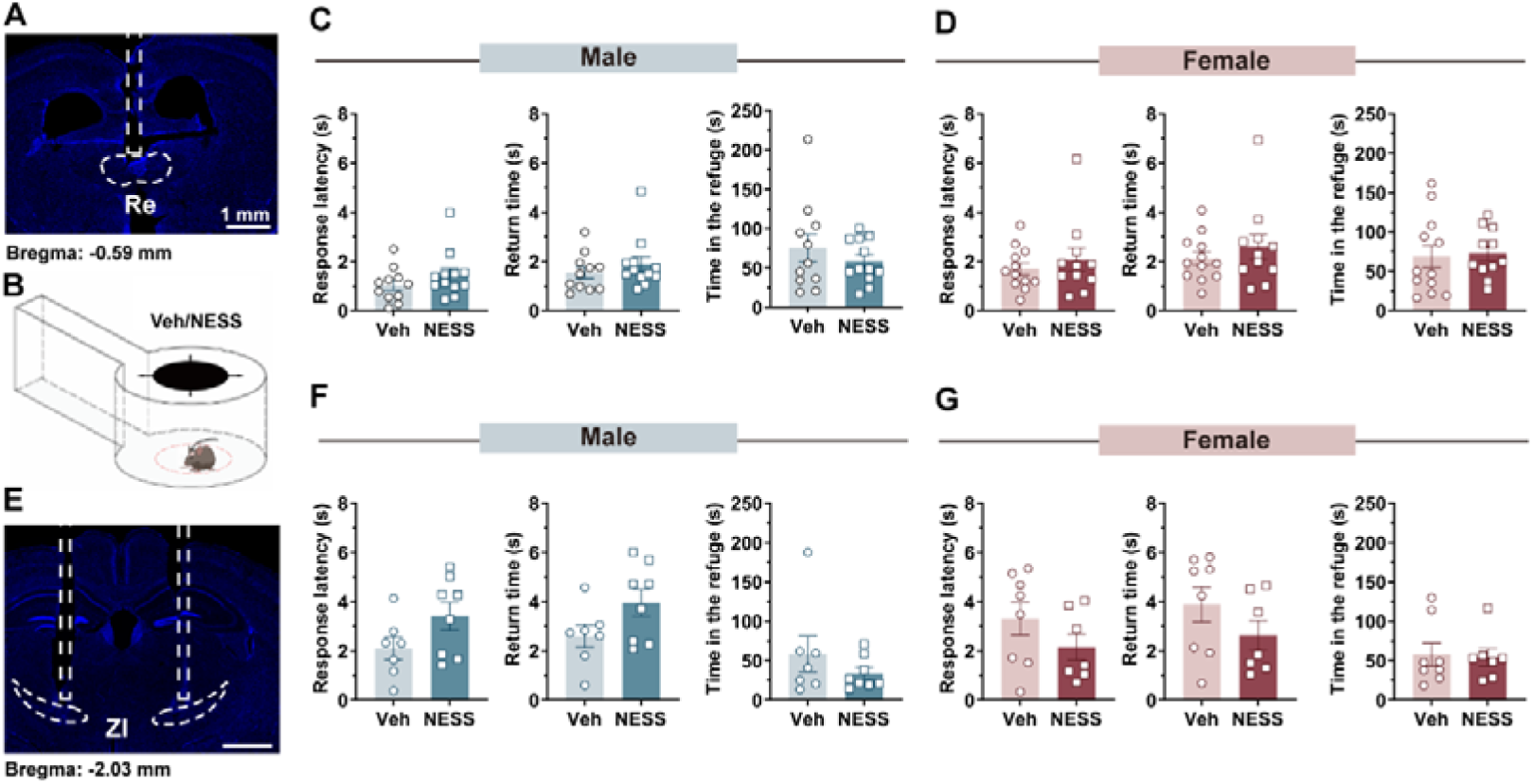
CB_1_R antagonism in Re or ZI lacks effect on defensive behavior in either sex, related to Figure 3. (A) Representative image showing the track cannula placement in the reuniens thalamic nucleus (Re). (B) Schematic representation of the looming stimuli paradigm conducted in a refuge-containing open field apparatus. (C and D) Metrics of defensive behavior (from left to right: response latency, return time, and time in the refuge after looming) for male (C) and female mice (D) in the vehicle (Veh) and NESS 0327 (NESS) groups. (E) Representative image showing the track cannula in the zona incerta (ZI). (F and G) Defensive behavior metrics (from left to right: response latency, return time, and time in the refuge after looming) for male (F) and female mice (G) in the Veh and NESS groups. Data are presented as mean ± SEM. n=11 male Veh group, 12 male NESS group (C); 12 female Veh group, 11 female NESS group (D); 7 male Veh group, 8 male NESS group (F); 8 female Veh group, 7 female NESS group (G). Unpaired Student’s t test with the Mann-Whitney test was performed. For detailed *p* values, refer to Table S1.

**Figure S12.**
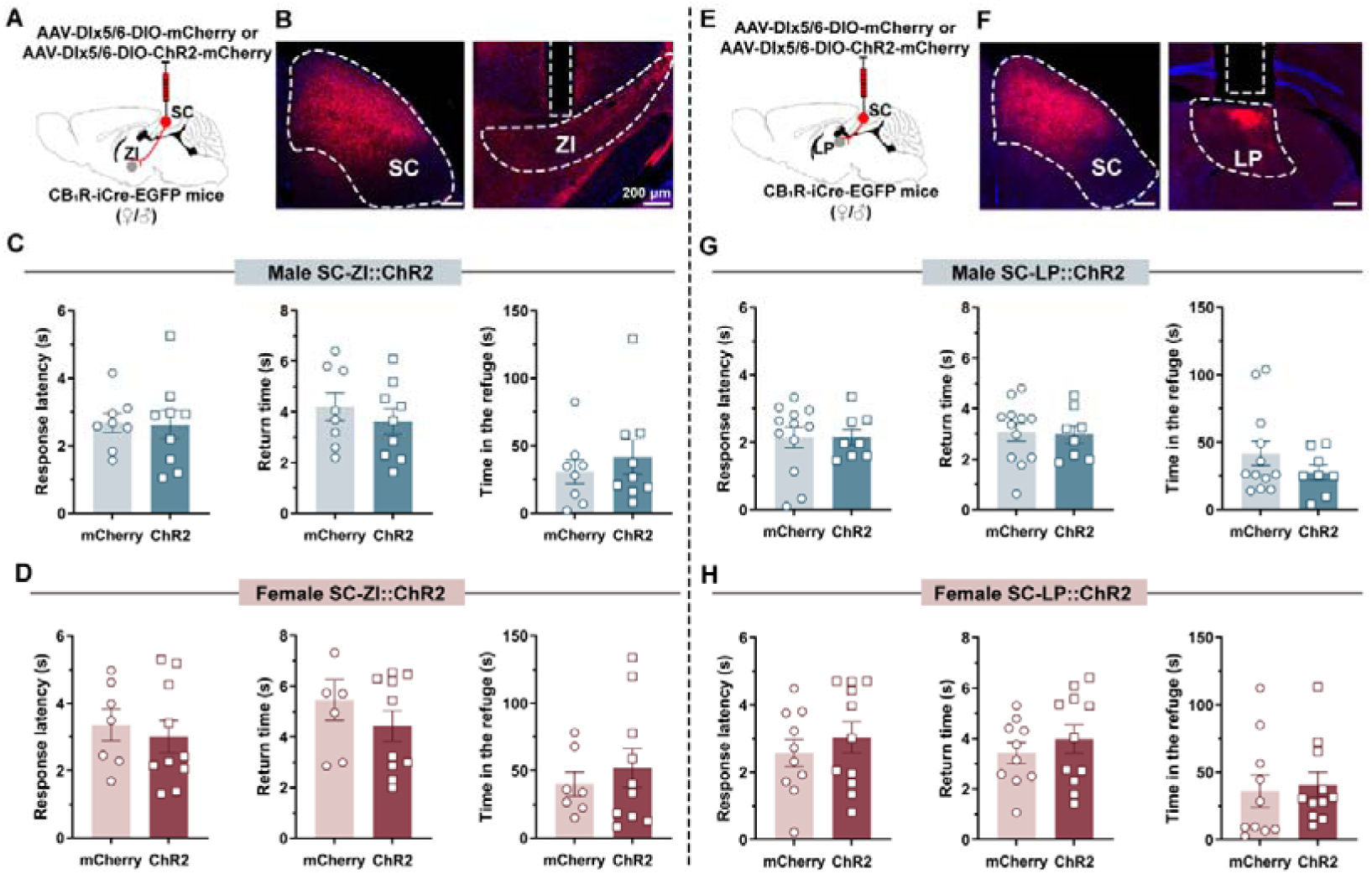
Activating CB_1_R^+^Dlx5/6^+^ SC-ZI/LP terminals lacks effect on defensive behavior in either sex, related to Figure 4. (A and B) Viral strategy and histology of CB_1_R-expressing SC-ZI Dlx5/6^+^ terminals. The dotted box highlights the optic fiber track. Scale bar, 200 μm. SC, superior colliculus; ZI, zonal incerta. (C and D) Statistic analysis of defensive behavior metrics, including time spent initiating a response, returning to the refuge, and remaining in the refuge after looming stimuli during opto-stimulation of SC-ZI^CB1R+ Dlx^^5/6+^ terminals in males (C) and females (D). (E and F) Viral strategy and histology analysis of the SC-LP^CB1R+Dlx^^5/6+^ terminals. The dotted box highlights the optic fiber track. Scale bar, 200 μm. LP, lateral posterior thalamus. (G and H) Statistic analysis of defensive behavior metrics, including the time spent initiating a response, returning to the refuge, and remaining in the refuge after looming stimuli during opto-stimulation of SC ^CB1R+Dlx^^5/6+^ axons in the LP for males (G) and females (H). Data are presented as mean ± SEM. n = 8 male mCherry group, 9 male ChR2 group (C); 7 female mCherry group, 10 female ChR2 group (D); 12 male mCherry group, 8 male ChR2 group (G), 10 female mCherry group, 11 female ChR2 group (H). Unpaired Student’s t test with the Mann-Whitney test was performed. For detailed *p* values, refer to Table S1.

**Figure S13.**
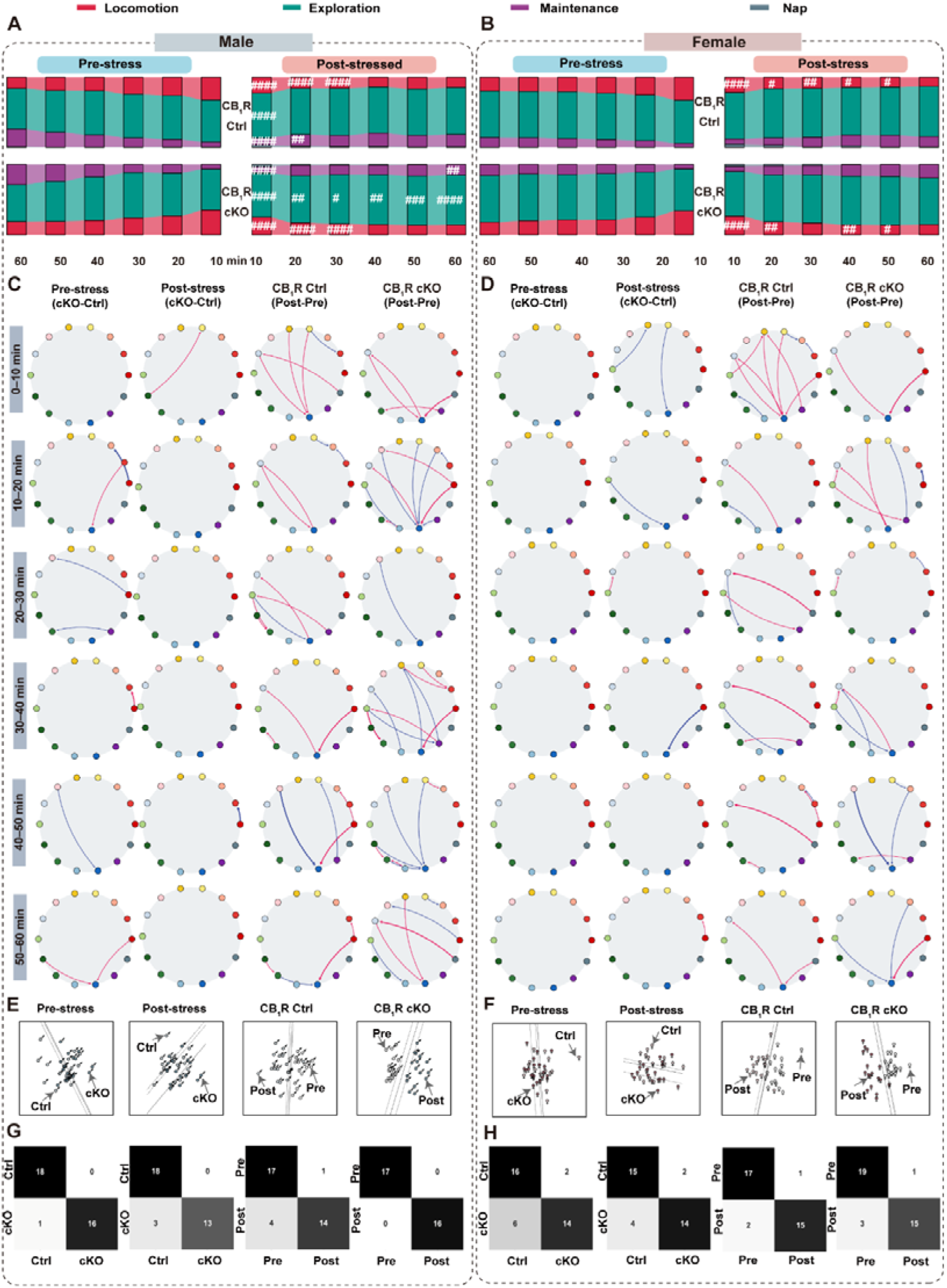
Chronic stress alerts spontaneous behavior dynamics, related to Figure 6. (A and B) Proportion of clusters during every 10-min epoch for male (A) and female (B) CB_1_R Ctrl (top) and CB_1_R cKO (bottom) groups. Data are presented as mean ± SEM. Male CB_1_R Ctrl (n=18 pre-stress, 17 post-stress); Male CB_1_R cKO (n=18 pre-stress, 16 post-stress); Female CB_1_R Ctrl (n=17 pre-stress, 20 post-stress); Female CB_1_R cKO (n=17 pre-stress, 18 post-stress). #*p*<0.05, ##*p* < 0.01, ###*p* < 0.001, ####*p* < 0.001 post-stress versus pre-stress in corresponding group. Two-Way ANOVA with Sidak’s multiple comparisons post hoc test was performed. For detailed *p* values, refer to Table S1. (C and D) Comparison of movement transition probabilities for males (C) and females (D). From left to right, pre-stress (CB_1_R cKO minus CB_1_R Ctrl), post-stress (CB_1_R cKO minus CB_1_R Ctrl), CB_1_R Ctrl group (post-stress minus pre-stress), CB_1_R cKO group (post-stress minus pre-stress). Each movement bout is represented as a state. (E and F) PCA visualization of behavioral features for males (G) and females (H) across different groups. (G and H) Random forest classifier trained on 14-dimensional movement fractions and frequency to predict group labels for males (G) and females (H) across different groups.

**Figure S14.**
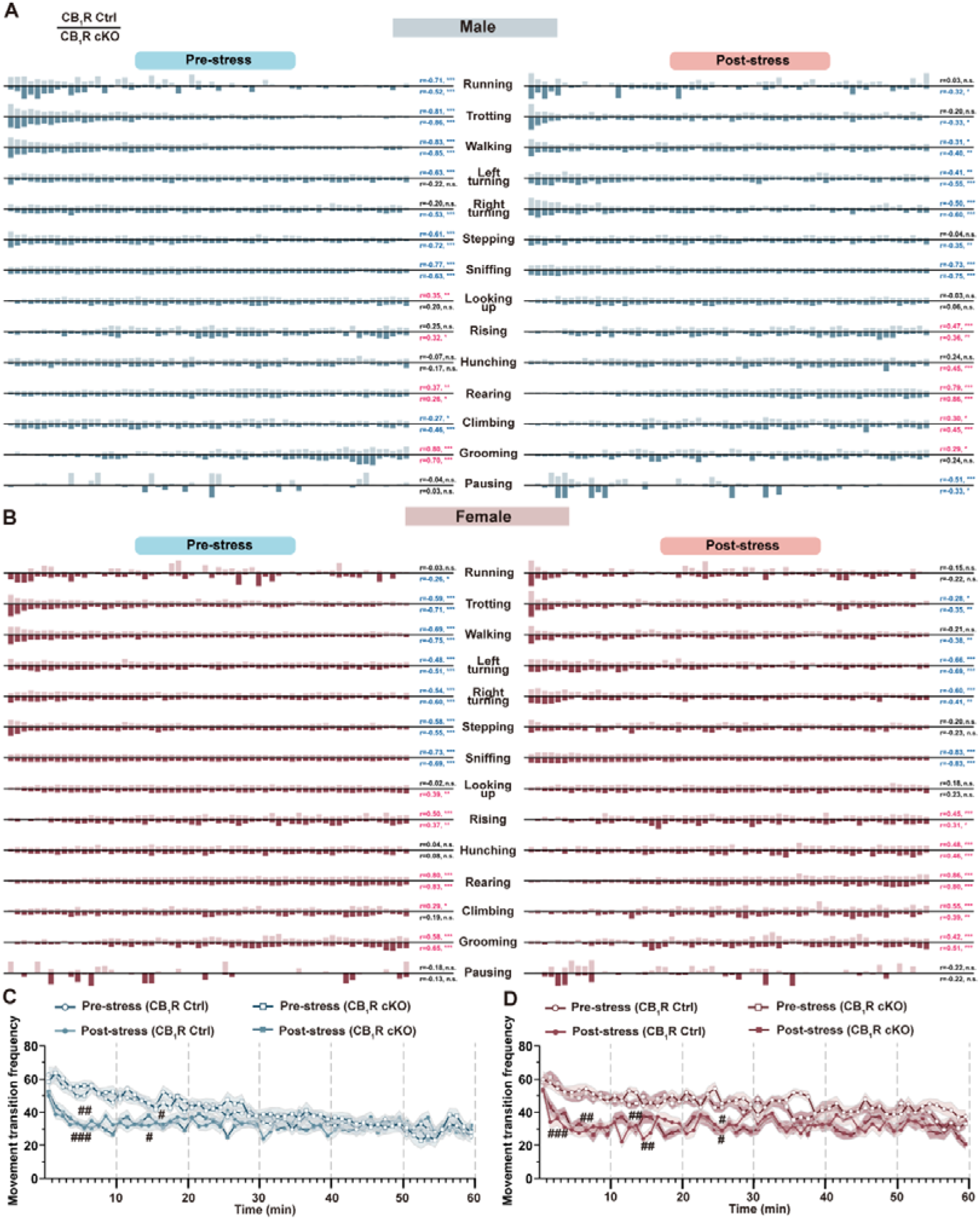
Dynamics of spontaneous behavior over 60 min period, related to Figure 6. (A and B) Temporal fraction of each movement during pre- and post-stress stages for males (A) and females (B) in CB_1_R Ctrl (top) and CB_1_R cKO (bottom) groups. Data are presented as the mean normalized values over the time course for each movement. (C and D) Movement transition frequency during pre- and post-stress stages for males (C) and females (D) in CB_1_R Ctrl and CB_1_R cKO groups. Data are presented as mean ± SEM. Sample sizes are consistent with those in Figure S13. #*p*<0.05; ##*p*<0.01; ###*p*<0.001 post-stress versus pre-stress in corresponding group. Two-Way ANOVA with Sidak’s multiple comparisons post hoc test was performed. For detailed *p* values, refer to Table S1.

**Table S3.** Parameters in the Dlx5/6 experiments during looming stimuli, related to Figure 2.

| Type | Sex | Group | Speed<br>(mm/s)<br>Mean±SEM | Nose height<br>(mm)<br>Mean±SEM | Back height<br>(mm)<br>Mean±SEM |
| --- | --- | --- | --- | --- | --- |
| Flight | Male | Ctrl (n=22) | 153.4±10.49 | 21.09±2.313 | 25.21±1.049 |
|  |  | cKO (n=22) | 149±11.72 | 16.02±2.975 | 26.57±0.8789 |
|  | Female | Ctrl (n=16) | 133.9±17.23 | 21.09±2.313 | 25.26±0.9304 |
|  |  | cKO (n=16) | 166.2±9.934 | 16.02±2.975 | 23.68±1.177 |
| Non-Flight | Male | Ctrl (n=6) | 52.09±18.88 | 27.39±7.068 | 28.7±1.113 |
|  |  | cKO (n=2) | 96±9.378 | 22.38±6.334 | 26.71±3.041 |
|  | Female | Ctrl (n=4) | 62.47±10.22 | 27.42±4.827 | 27.07±1.087 |
|  |  | cKO (n=5) | 81.16±23.89 | 27.41±7.222 | 28.71±2.119 |

**Table S4.** Definition of movements, related to Figure 2.

| **Movement** | **Definition** |
| --- | --- |
| Flight | Mouse locomotes with relatively high speed following looming stimuli |
| Running | Mouse locomotes with relatively high speed |
| Trotting | Mouse locomotes with a slow and intermittent pace |
| Walking | Mouse locomotes with relatively low speed |
| Left turning | Mouse bends the body to the left or turning body to the left while walking |
| Right turning | Mouse bends the body to the right or turning the body to the right while walking |
| Stepping | Mouse takes a step forward with a short distance |
| Rotating | Mouse turns around with low speed and moves a short distance |
| Sniffing | Mouse twitches its nose to explore the environment at a relatively low speed |
| Looking up | Mouse Looks up with four legs on the ground |
| Rising | Mouse lifts its front legs but doesn't stand up |
| Hunching | Mouse stands on its hind legs while the dorsal is bent |
| Rearing | Mouse stands on hind legs with straight back |
| Grooming | Mouse licks, grooms its fur with the forepaws, or scratches its fur using any limb |
| Pausing | Mouse is still, crouches, or lies down on the floor without obvious horizontal displacement |
| Freezing | Mouse is completely absence of movement |

**Table S5.** Parameters in the CaMKIIα experiments during looming stimuli, related to Figure 2.

| Type | Sex | Group | Speed<br>(mm/s)<br>Mean±SEM | Nose<br>height<br>(mm)<br>Mean±SEM | Back height<br>(mm)<br>Mean±SEM |
| --- | --- | --- | --- | --- | --- |
| Flight | Male | Ctrl (n=26) | 150.2±9.45 | 15.33±2.036 | 22.54±0.8555 |
| | | cKO (n=15) | 151.5±12.17 | 15.4±2.073 | 23.23±1.099 |
| | Female | Ctrl (n=15) | 151.3±15.02 | 16.36±3.532 | 21.54±1.256 |
| | | cKO (n=8) | 142.4±16.97 | 16.34±2.449 | 22.53±1.21 |
| Non-Flight | Male | Ctrl (n=4) | 35.65±7.68 | 35.55±9.42 | 26.66±2.281 |
| | | cKO (n=12) | 48.26±4.861 | 35.15±4.261 | 29.94±1.479 |
|  | Female | Ctrl (n=0) |  |  |  |
| | | cKO (n=4) | 45.1±4.248 | 42.15±6.576 | 28.46±2.759 |

**Table S6.** Definition of movements and clusters, related to Figure 5, Figure 7.

| **Clusters** | **Movement** | **Definition** |
| --- | --- | --- |
| Locomotion | Running | Mouse locomotes with relatively high speed |
|  | Trotting | Mouse locomotes with a slow and intermittent pace |
|  | Walking | Mouse locomotes with relatively low speed |
|  | Left turning | Mouse bends the body to the left or turning body to the left while walking |
|  | Right turning | Mouse bends the body to the right or turning the body to the right while walking |
|  | Stepping | Mouse takes a step forward with a short distance |
| Exploration | Sniffing | Mouse twitches its nose to explore the environment at a relatively low speed |
|  | Looking up | Mouse Looks up with four legs on the ground |
|  | Rising | Mouse lifts its front legs but doesn't stand up |
|  | Hunching | Mouse stands on its hind legs while the dorsal is bent |
|  | Rearing | Mouse stands on hind legs with straight back |
|  | Jumping | Mouse jumps with its hind legs completely off the ground |
| Maintenance | Grooming | Mouse licks, grooms its fur with the forepaws, or scratches its fur using any limb |
| Nap | Pausing | Mouse is still, crouches, or lies down on the floor without obvious horizontal displacement |

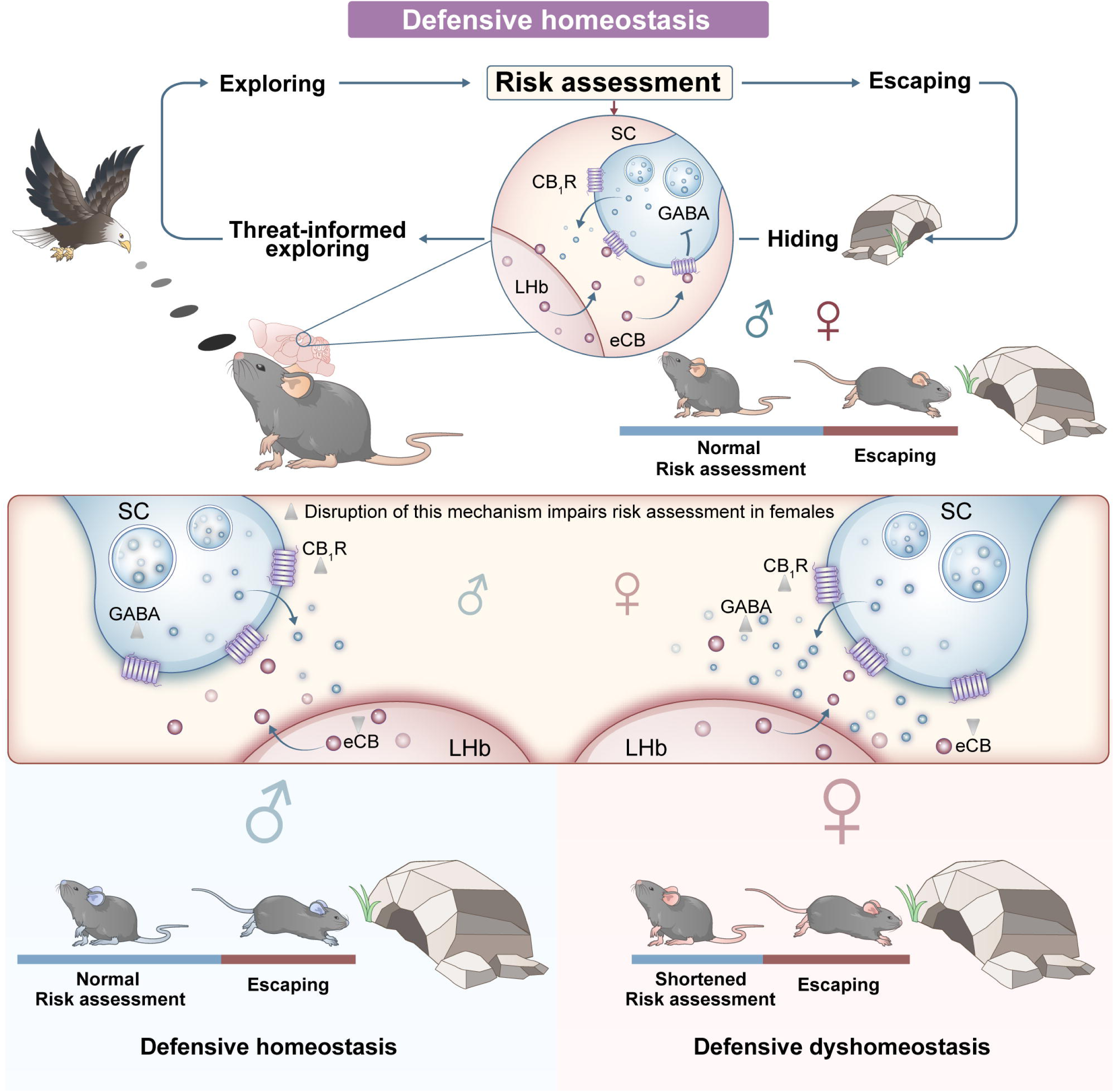

